# Rapidly Inferring Personalized Neurostimulation Parameters with Meta-Learning: A Case Study of Individualized Fiber Recruitment in Vagus Nerve Stimulation

**DOI:** 10.1101/2022.09.06.506839

**Authors:** Ximeng Mao, Yao-Chuan Chang, Stavros Zanos, Guillaume Lajoie

**Author notes:** Corresponding Author (G. Lajoie).

## Abstract

**Objective:** Neurostimulation is emerging as treatment for several diseases of the brain and peripheral organs. Due to variability arising from placement of stimulation devices, underlying neuroanatomy and physiological responses to stimulation, it is essential that neurostimulation protocols are personalized to maximize efficacy and safety. Building such personalized protocols would benefit from accumulated information in increasingly large datasets of other individuals’ responses.

**Approach:** To address that need, we propose a meta-learning family of algorithms to conduct few-shot optimization of key fitting parameters of physiological and neural responses in new individuals. While our method is agnostic to neurostimulation setting, here we demonstrate its effectiveness on the problem of physiological modeling of fiber recruitment during vagus nerve stimulation (VNS). Using data from acute VNS experiments, the mapping between amplitudes of stimulus-evoked compound action potentials (eCAPs) and physiological responses, such as heart rate and breathing interval modulation, is inferred.

**Main results:** Using additional synthetic data sets to complement experimental results, we demonstrate that our meta-learning framework is capable of directly modeling the physiology-eCAP relationship for individual subjects with much fewer individually queried data points than standard methods.

**Significance:** Our meta-learning framework is general and can be adapted to many input-response neurostimulation mapping problems. Moreover, this method leverages information from growing data sets of past patients, as a treatment is deployed. It can also be combined with several model types, including regression, Gaussian processes with Bayesian optimization, and beyond.

## 1. Introduction

**Table 1:**
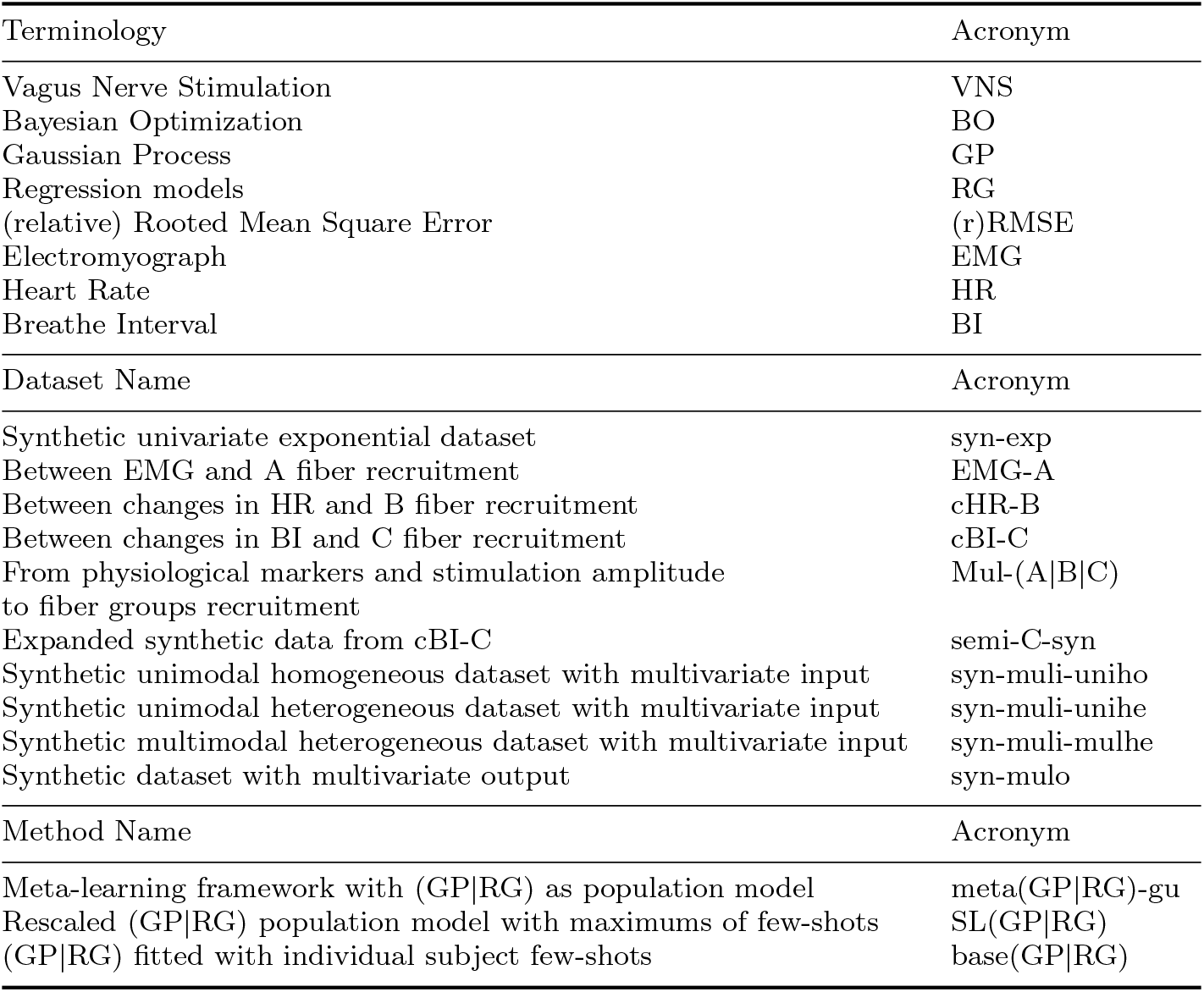
Table of Key Acronyms in the Manuscript.

Neurostimulation is growing in popularity as a potential treatment for several ailments, for example, spinal cord stimulation (SCS) for chronic pain [1], deep brain stimulation (DBS) for Parkinson’s disease [2] and vagus nerve stimulation (VNS) for epilepsy [3]. It is administered via either invasive or non-invasive approaches to deliver stimulation to nervous system [4]. This can be done either in the central or peripheral nervous systems for varied targeted outcomes. While electrical stimulation is widely implemented for neurostimulation, other modalities exist such as ultrasound [5]. An important part of developing neurostimulation treatments is to accurately estimate a subject’s neural and physiological response map to stimulation, which can then be used for personalized optimization. Moreover, in more generality, deriving the personalized mapping between neural activity (evoked or not) and physiological response can be of great value. However, this is challenging even in the presence of ever growing datasets from past patients, mainly because of inherent variability in subject to subject responses, but also because of varying implant placement and other physiological factors [7]. It is therefore important to leverage this acquired data from the past, while avoiding the pitfalls of simply fitting and using population mean response models. Data from the current patient is also crucial in learning an accurate personalized model, but the amount of queries and the intensity range must often be constrained in practice, for safety and other practical concerns. To address this problem, we propose a flexible and robust meta-learning framework to perform *few-shots* learning of key response fitting parameters given a new subject and a pool of past model fits data.

Few-shot learning [8,9] is a widely studied learning paradigm in machine learning, where the learning model is asked to solve each task while observing only a small fractions of the data. In the context of neurostimulation, each shot can be a query to the subject’s response map. Therefore, the ability to do well in few-shot learning is beneficial for neurostimulation, considering both that it can reduce the number of stimulations for calibration purposes, and that it can avoid the amount of time required to process the additional signals. Attempting to generalize well with only a small training set can be challenging, but it can be facilitated by leveraging the experience of learning other tasks that are similar, which is the core concept of meta-learning [10, 11], also known as “learning to learn”. The overarching training objective of few-shot learning, solved by meta-learning approaches, focuses on improving a model’s ability to adapt and generalize with access to only few samples from each task, while leveraging meta learned prior knowledge from previously available tasks. Note that in meta-learning, we have data both from other tasks as well as the few-shots from the current task. Therefore, to better distinguish between the two sources of information, the former is commonly termed as the meta-training set.

By viewing the personalization of one single subject as one task, we can draw direct parallel to meta-learning, where the goal of meta-learning becomes to extract, from a population of past subjects, as much information that is useful for a personalized model of a new subject to be optimized in open or closed loops settings. To exemplify the use of our proposed meta-learning framework, in this manuscript, we explore the physiological modeling of a new individual’s fiber recruitment in VNS and the associated physiological responses such as heart rate and breathing interval modulation. We conduct our analysis based on the published data found in [21]. Here, we focus on modelling the mapping between physiological observables and neural activity evoked by neurostimulation, and we extract a model of this mapping from very few sampled pairs of neural activity and physiological signals (few shots).

VNS is an emerging treatment for a multitude of disorders [3, 6, 13–17] including epilepsy, depression, anxiety, Alzheimer’s disease, chronic pain and rheumatoid arthritis. It stimulates the vagus nerve, a main nerve of peripheral nervous systems that extends from the brainstem to the abdomen and is responsible for vast essential body functions, such as heart rate, breathing and digestion [12]. Accumulating evidence suggests that varying the stimulation parameters for individual subjects can improve the efficacy of VNS [18,19]. Hence, to develop targeted and personalized clinical interventions, it is crucial to acquire quantifiable fiber recruitment during VNS on a single subject basis. Obviously, this requirement is shared with several other stimulation modalities and observables, which makes the particular problem a representative case study to evaluate meta-learning.

Based on the known correlations between certain physiological markers and specific fiber groups [22, 23], one way to obtain the fiber responses’ amplitudes is through modeling the physiological effects in response to the neurostimulation. Note that this is alternative to obtaining the fiber responses directly from neural recording which is difficult in clinical applications, and that it is much easier to obtain the physiological readings through non-invasive ways. Recent work by Chang *et al* [21] demonstrated linear or non-linear relationships in a rat model for each of the three pairs of physiological markers and fiber amplitudes of stimulus-evoked compound action potentials (eCAPs): between cervical electromyography (EMG) and A-fiber, between changes in heart rate (∆HR) and B-fiber and between changes in breathing interval (∆BI) and C-fiber. The authors further derived and validated regression models for each fiber group from either univariate and multivariate inputs, given normalized responses from each individual. However, due to the needs of working with subject-dependent normalization factors, the trained regression models cannot be applied directly to unseen subjects, which is a hindrance for these normalized response models to be used in deployed VNS treatments.

In this study, we provide a use case of our proposed meta-learning framework to this problem, where the individual subject’s operating ranges of both physiological and fiber responses’ amplitudes are the key response fitting parameters to be learned. More specifically, the framework is tasked with both the learning of a population model from a pool of past subjects and the inference of the factors of normalization given the few shots from a new subject. Combining the two pieces of information, a personalized prediction model can then be derived for the new individual, using as few query points or “shots” (stimulation with distinct parameter values) as possible. With prior knowledge learned from available meta-training data, our framework creates an average model (hereon referred to as population model), and then adapts it accordingly for each new subject given few shots of the measurements. The framework is by design compatible with any model type as population model, creating a family of algorithms, and we investigate two of them in our implementations: regression and Gaussian process (GP) [24]. We validate the performance of our proposed metalearning framework in multiple evaluations on both publicly available experimental data and synthetically created artificial data, and we find accurate and consistent performance across all evaluations regardless of the choice of population models.

The main contributions of our manuscript are summarized as the followings:

- We propose the use of meta-learning as a promising and generic solution to the challenge of personalization in neurostimulation.
- As a concrete case study, we propose a meta-learning framework for physiological modeling of fiber recruitment in VNS, capable of deriving individual subjects’ models using only few data points, by leveraging the extracted prior knowledge from the population.
- We provide evaluations on both synthetic and publicly available experimental data, with both univariate and multivariate relationships. The results show the framework’s competitive performance compared to other standard methods, across multiple scenarios, especially when the number of few-shots is small and when the accessible few-shots are limited within a subset of the data space.

Importantly, we reiterate that our proposed framework is general and can be exploited in various settings, as we discuss later on.

The manuscript is structured as follows. In Section 2, we illustrate the process of utilizing the meta-learning framework to predict individual subject’s fiber responses’ amplitudes, and we show the results of evaluations on both real experimental and synthetic data, using the framework. We then provide a discussion where we outline the limitation and future work in Section 3. Lastly in Section 4, we discuss the relevant topics of the meta-learning framework, including the problem formulation of few-shot learning, the related works, the background knowledge of several techniques used in the framework, and the details regarding the utilized synthetic and real experimental data.

## 2. Results

### 2.1 Meta-Learning Framework for Few-shot Prediction of Individual Subjects’ Fiber Responses’ Amplitudes

In Figure 1(a), we provide an illustration on how to use the proposed meta-learning framework to conduct few-shot prediction on individual subjects’ fiber amplitudes, and the diagram of the framework is shown in Figure 1(b). Here we assume univariate function relationship, and multivariate case is analogous. Pseudo-code of the framework is presented in Algorithm 1. Acting as the backbone for all the future individual models, the framework compiles the population model a priori, using the normalized data from all subjects in the meta-training set, which in our case is more fitting to refer as the population set. Note that we will use meta-training set and population set interchangeably for the rest of this manuscript. The population model is supposed to reflect the prior knowledge extracted from the past data, embodied here as a population mean fit. There is no restriction on the model type used as population model, with notable candidates as GP and regression models. For readers familiar with meta-learning, the parameters of the population model can be viewed as the meta parameters, considering they are learned from the pool of population and serve as the extracted prior knowledge.

**Figure 1.**
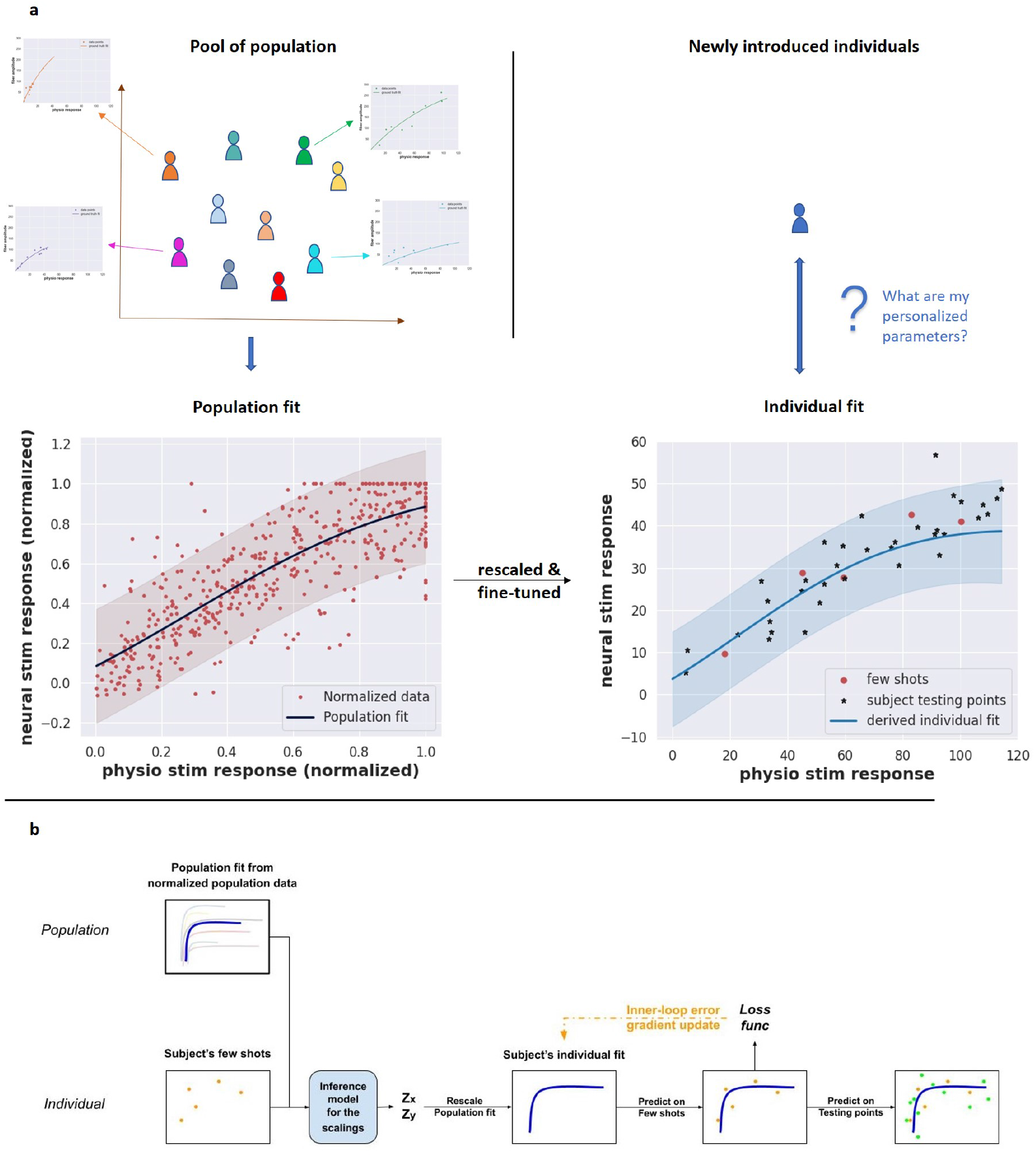
The proposed meta-learning framework to predict individual subject’s fiber responses’ amplitudes. (a) Illustrations on utilizing the meta-learning framework to predict on newly introduced subject, where the objective is to learn a mapping between individual subjects’ neural and physiological responses’ amplitudes in response to stimulation. (b) Diagram of proposed meta-learning framework.

For each subject in both meta-training and meta-testing set, the framework takes few shots (queries) from it and infers its scaling factors (*z*_*x*_, *z*_*y*_), where *x* and *y* correspond to physiological and fiber responses’ amplitudes respectively. Finally the subject’s individual model, or individual fit in this case, is derived by rescaling the population fit accordingly. Our framework uses Bayesian optimization (BO) [35] as the inference model, aiming to minimize the prediction error on the few-shots by the rescaled population fit, with respect to the inferred scaling factors. Therefore, we expect the resulted scaling factors to be proper estimates of the operating ranges of both physiological and fiber responses’ amplitudes. The resulted individual model can then be evaluated on the unseen data points of the same subject, which will be denoted as subject test points. Generalization performance of the framework will be conducted on meta-testing subjects.

The scaling operation alone might not be sufficient for personalization, and to cope with that, we incorporate the inner-loop error feedback into the framework, as an one-step gradient update based on the feedback from the few-shots. The gradient-based adaptation is conducted on the parameters of each individual model, initially copied from the parameters of the population model. It allows for more expressive tuning of the individual fit. In Figure 1(b), the inner-loop error feedback path is illustrated as yellow dashed lines. Besides feedback on the few-shots, feedback on the subject test points can also be used during meta-training, which we analogously refer to as outer-loop error feedback. The outer-loop error can be used to improve the overall few-shot learning procedure. Here, for all the evaluations to be described in the following sections, we used the outer-loop error only to guide the search of the learning rate of the inner-loop gradient update, in a separate training process beforehand, and we fixed the resulted learning rate during the actual evaluation. Since the outer-loop feedback path is active only in the pretraining phase, we exclude that from the diagram in Figure 1(b).

#### Algorithm 1 Few-shot Prediction on Individual Fiber Amplitudes from Physiological Responses

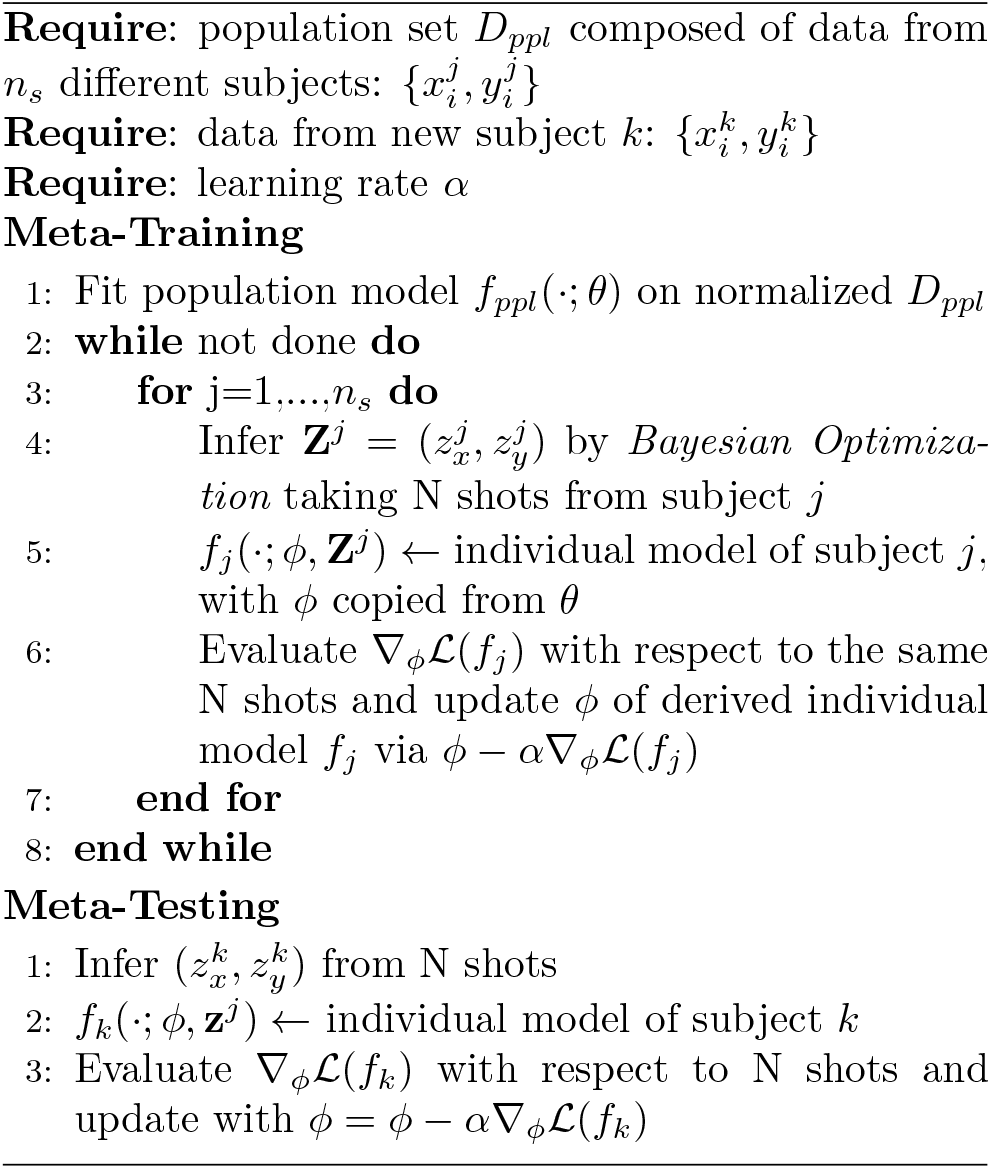

#### 2.1.1 Variants of algorithms under evaluations

By varying the choices of population models, our meta-learning framework defines a family of algorithms. In evaluating our approach, we consider two variants: `metaRG-gu` and `metaGP-gu` with either regression or GP as the population models (here `gu` means inner-loop gradient update). Whenever available, we allowed those with regression population model (`RG`) to use the best possible function forms during evaluations. For example, closed-form exponential function will be used if we know a priori the data demonstrates an exponential relationship. On the other hand, for GP, we used a generic Matern kernel for most of the evaluations and a linear kernel for data points complying a linear relationship.

In particular, a special case of our framework is functionally similar to standard supervised learning with normalized data. Here we also consider a naive version of the framework with a trivial inference model using the maximums of the few-shots for normalization, and without the inner-loop gradient update. We refers to them as `SLRG` and `SLGP` highlighting their connections to standard supervised learning. The details of these methods are discussed in next subsection. Note that although they are in principle variants of our framework, in this manuscript we treat them primarily as baselines for comparison with non-trivial meta-learning models.

### 2.2 Evaluation Metrics and Baseline Comparisons

Two evaluation metrics are used for performance comparison. The first is the test rooted mean squared error (RMSE), which is the prediction error of derived individual model on subject test points.

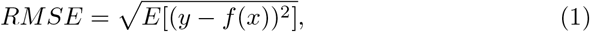

or the normalized version of it, relative RMSE (rRMSE), where the predictions were normalized by ground truth scaling factors.

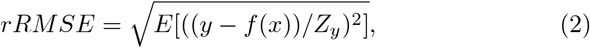

In this manuscript, for both RMSE and rRMSE metrics, we report for each datasets their per-subject mean and per-subject standard deviation divided by the square root of the number of subjects, where per-subject standard deviation is calculated by averaging of all subjects’ standard deviations across different trials. We consider three sets of baselines in the evaluations. The first set of baselines includes `SLRG` and `SLGP`, which stand for supervised learning with either the population models. In our case, the supervised learning paradigm would mean to train on the normalized data points from subjects in meta-training set and evaluate on the normalized data points from those in meta-testing set, as was done in previous publications [21]. Adopting that paradigm in the few-shot learning setup requires extra steps of normalizing the meta-testing few-shots to feed in the model and scaling the output back for proper comparisons. Here the normalization was done simply with the maximum of its few-shots. As mentioned above, this set of baselines can also be viewed as naive variants of our meta-learning framework without BO nor the inner-loop gradient update. The second set includes `baseRG` and `baseGP`, which are regression and GP fitted directly on each subject’ few-shots, respectively, and no forms of normalizations are conducted in this set of baselines. Note that although the closed form of regression model in `baseRG` is still coming from prior knowledge, this set of baselines represents the case where prior knowledge on population fits’ shape is not explicitly utilized in deriving the individual fits. The last set is a popular deep learning based method called model agnostic meta-learning (MAML) [11], and it is an optimization-based meta-learning method that aims to learn a good initialization of the model parameters, as the extracted prior knowledge and the meta parameters, with inner-loop gradient update to adapt to each individual task. This baseline represents a generic meta-learning solution without tailoring to the specialities of our problem.

### 2.3 Performance Evaluations

Four sets families of datasets are tested to examine the performance of our proposed meta-learning framework. For the first one, we built datasets from scratch with purely synthetic univariate subjects, generated from one-dimensional (1-d) exponential functions and noise. With this evaluation, we aim to verify the performance of our meta-learning framework in the ideal situation where all subjects are exactly rescaled versions of the same base function. Second, to investigate the framework’s performance in practical settings, we conducted evaluations on publicly available real experimental data from previous publication [21], with either univariate or multivariate inputs.

Given the importance of the population model in our meta-learning framework, it is worthwhile to investigate how various aspects of population set could affect the few-shot performance. Therefore, we synthetically expanded one real univariate dataset as the third set of datasets to evaluate. We term this set of datasets semi-synthetic, highlighting its connections to the real data. Lastly, to investigate the scaling performance of our framework to higher dimensional spaces, we tested it on synthetic datasets with multivariate inputs or outputs. For the details descriptions and results of this last evaluation, please refer to Appendix C.

#### 2.3.1 Evaluations on purely synthetic univariate datasets

Figure 2 shows the results on one purely synthetic dataset where all subjects are scaled versions of one exponential function. We refer to this dataset as `syn-exp`, and refer the reader to Section 4.5 for details regarding subject generation procedure and the specifications of the used exponential function. We performed leave-one-out cross-validations and reported the average rRMSE per number of shots (i.e. random stimulation query) in Figure 2(b) and (c). These sub-figures show the competitive performance of our proposed meta-learning framework, especially when the number of shots is small, meaning rapid inference of a model fit with very few sample points. We include also in the two sub-figures the red dashed line as the *dataset noise*, calculated as the averaged standard deviation of ground truth zero-mean noise for `syn-exp.` We can clearly observe the trends of all methods coverging to the dataset noise when more shots of the subject are made available, but both `metaGP-gu` and `metaRG-gu` approach it much earlier, with as small as 3 or 5 shots. On one hand, this demonstrates the power of utilizing the extracted prior knowledge from the population set, in relieving and even overcoming the lack of necessary information caused by only accessing to a small amount of data. On the other hand, as simple and straight-forward as the rescaling operation, when used within an appropriate framework of prior knowledge and inference model, can still outperform successful deep learning method like `MAML`. This highlights the importance of tailoring to the problem at hands. Furthermore, to investigate the role of BO, we report the performance comparison with or without the inner-loop gradient update in Appendix D. Although adding BO didn’t change much for this particular dataset, it did prove to be useful overall, especially when paired with GP population model.

**Figure 2.**
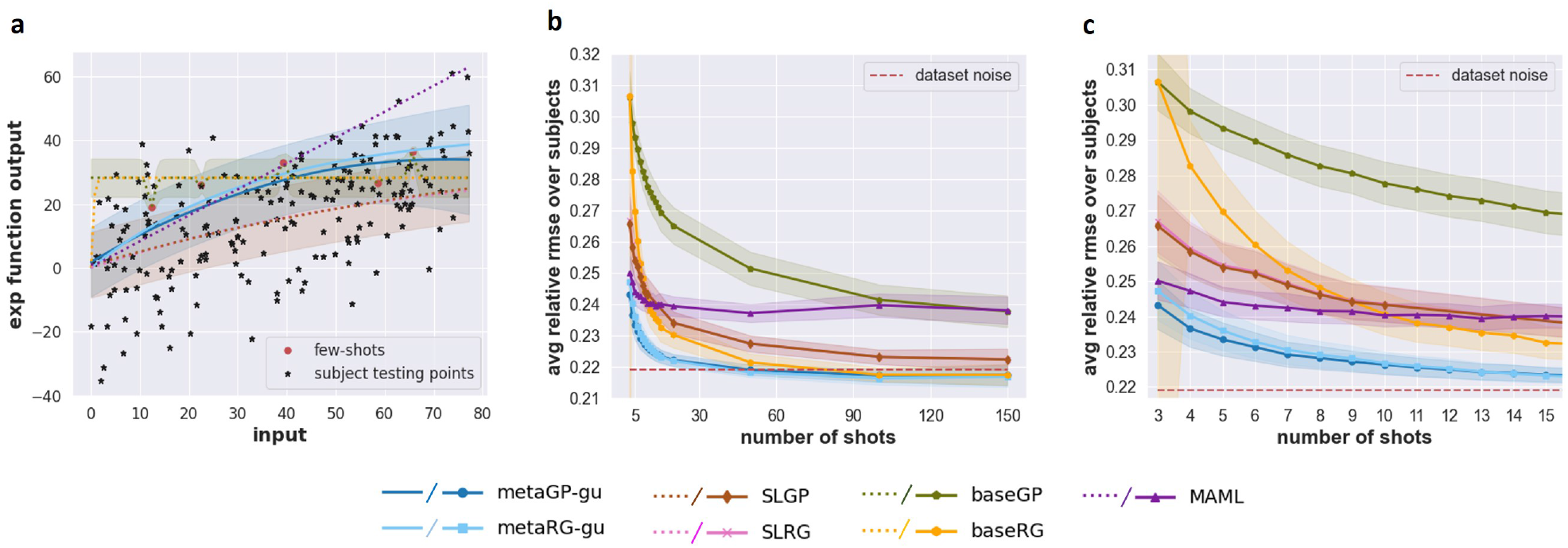
Performance comparison between the proposed meta-learning framework and the baselines on a purely synthetic dataset built from exponential function `syn-exp`. Note that the shaded areas in the sub-figures have different meaning. (a) One example of derived individual fits on 5-shots predictions. Shaded area shows prediction standard deviation for methods based on GP. (b) Performance curve of rRMSE with respect to the number of shots. Shaded area shows standard deviation of averaged performance per subject, divided by square root of number of subjects. (c) the same rRMSE curve as (b) but zooming in the region when number of shots is small (from 5 to 20).

In addition, this evaluation reveals the compatibility of our framework with GP. Despite being a powerful and generic method, GP alone, as shown with `baseGP` in Figure 2, might not be a good choice for fitting few-shot individual model for our problem. As shown in Figure 2(a), where sample individual fits on 5-shot predictions are plotted, `baseGP` struggles to recover the true exponential form and overfits to the noisy few-shots. However, a significant improvement can be observed in both the plotted curve and the reported rRMSE, when combining GP with our meta-learning framework. This justifies the flexibility of our framework regarding the choice of the population model, but more importantly, by being able to work with a generic method, this opens up the path of our framework to work with more datasets, for example, those not generated or well-described by a closed-form function.

#### 2.3.2 Evaluations on vagus nerve fiber recruitment datasets

We used the vagus nerve fiber recruitment data from [21] as experimental datasets to evaluate our method. The dataset includes univariate inputs, describing three 1-d physio-fiber relationships for VNS: from EMG to A-fiber, from ∆*HR* to B-fiber and from ∆*BI* to C-fiber, which we refer to as `EMG-A`, `cHR-B` and `cBI-C` respectively, as well as datasets of multivariate inputs, from the stimulation amplitudes and the three physiological markers to each of the fiber responses, which we refer to as `Mul-A`, `Mul-B` and `Mul-C` respectively.

To better compare with the initial study, we focus on modelling the relationship between physiological observables and the direct fiber activations instead of the stimulation amplitude used to evoke these activations, though our approach could easily be ported to this setting. The relationship between stimulation amplitude and fiber recruitment is straightforward and is described in [21].

Following the same procedure as in last section, leave-one-out cross-validations were adopted for performance comparisons, however here, we used RMSE instead as no ground truth scaling factors were available. Limited by experimental sample size, we evaluated on the univariate datasets with the number of shots between 2 and 4, and for the multivariate datasets with 3 and 5 shots. For regression models, we adopted the same model forms as the original paper, i.e. closed-form exponential and linear models for the univariate datasets and second degree polynomial regression models for the multivariate datasets. The closed-form regression models were used in both `SLRG` and `baseRG` baselines on the three real univariate datasets. As for the polynomial regression model, it was used in a baseline of a similar fashion as `SLRG`, but the normalizations on the inputs were applied on only one of the physiological markers: EMG, following the original paper. Due to this slight difference in rescaling schedule, we will refer to this polynomial regression baseline as `PolyRG` on the three real multivariate datasets. See Section 4.6 for more detailed descriptions on experimental data and the corresponding regression models.

Figure 3 shows our meta-learning methods univariate fits on held out subjects for this dataset. Comparative evaluation results are shown in Figure 4 and Figure 5, for real univariate and multivariate datasets respectively. We refer to Appendix B for the full results on all real experimental datasets in tables. Despite the limitations on the number of shots, we can observe the similar trends as in the last section. For univariate data, `baseGP` and `MAML` are the two worst-performance methods among the baselines, but likely distinct in their main causes. For `baseGP` it is likely due to the noise in the data points, which made the fitted GP aribitrary. As for `MAML`, the sizes of both the subjects and the data points in each subject were likely insufficient for a deep learning model, so we only include its results here for the sake of completeness. Same as the synthetic evaluations, `baseRG` suffers from unstableness when the number of shots is extremely small, but here the situation was even more severe as the large gap of degradation can also be observed in the RMSE mean, as shown in 2-shots results on `cBI-C`. Since `metaRG-gu` doesn’t display such performance drop in the evaluations, a probable explanation is that the shape of population fit provides a regularizing effect, preventing the individual fit to overfit to the noisy 2 data points. Thus, although it recovers quickly with more shots coming in, we can say that `baseRG` generally lacks of robustness to extremely small shots compared to our proposed meta-learning framework. Overall, the regression based methods do have a better performance than their GP counterparts, and one possible explanation is that exact function forms (exponential or linear) used in there are also a form of prior knowledge, coming from external expert knowledge. But we can also observe that the performance of `metaGP-gu` is on par with that of `metaRG-gu`, showcasing the capability of our meta-learning framework in closing the gap, when fitting the real data.

**Figure 3.**
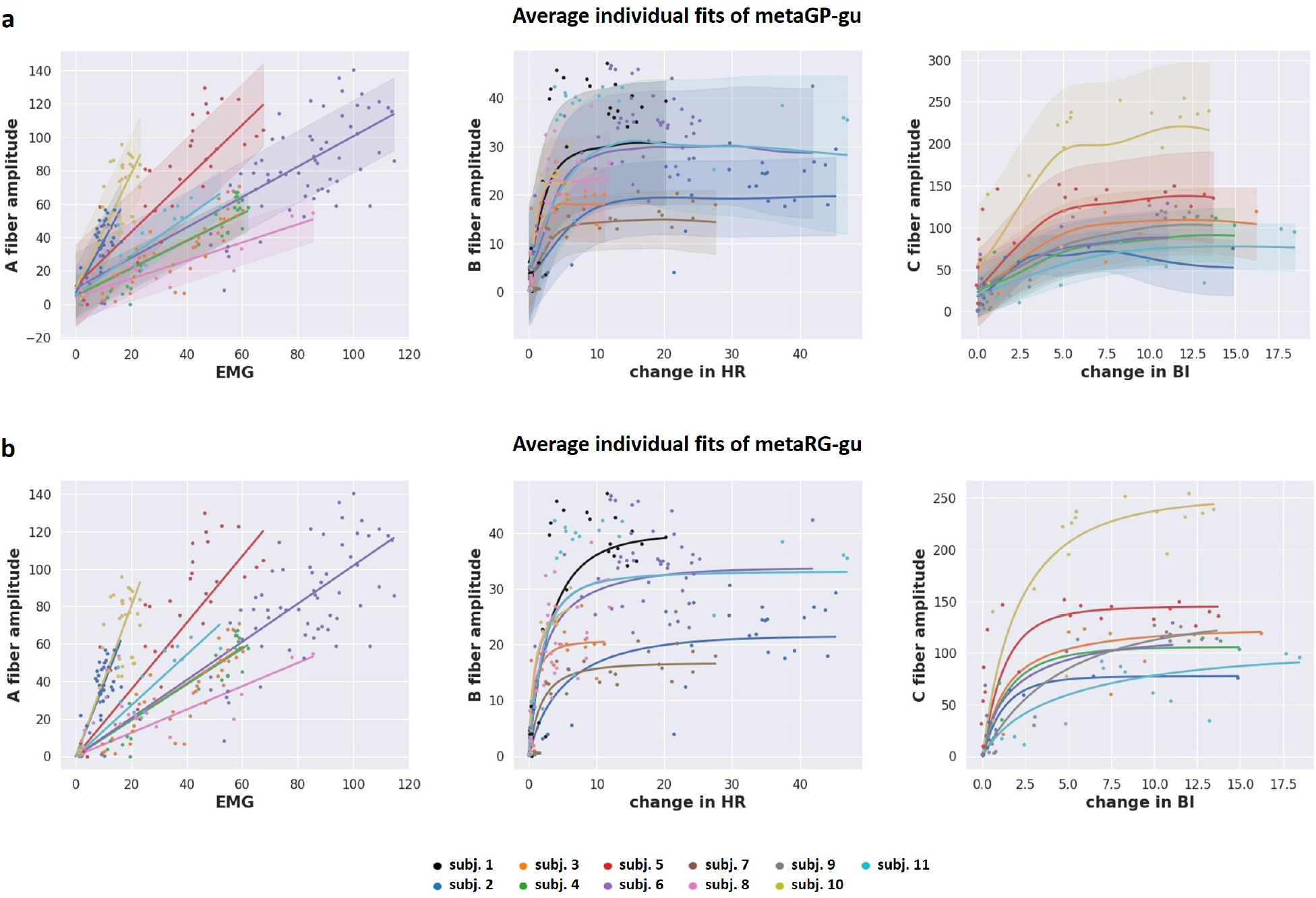
Derived individual fits on 3-shots predictions for all subjects in all three real univariate datasets, from (a) `metaGP-gu` and (b) `metaRG-gu`. Plotted are average mean over 50 repetitions, and average standard deviation on the GP prediction is shown for `metaGP-gu` only.

**Figure 4.**
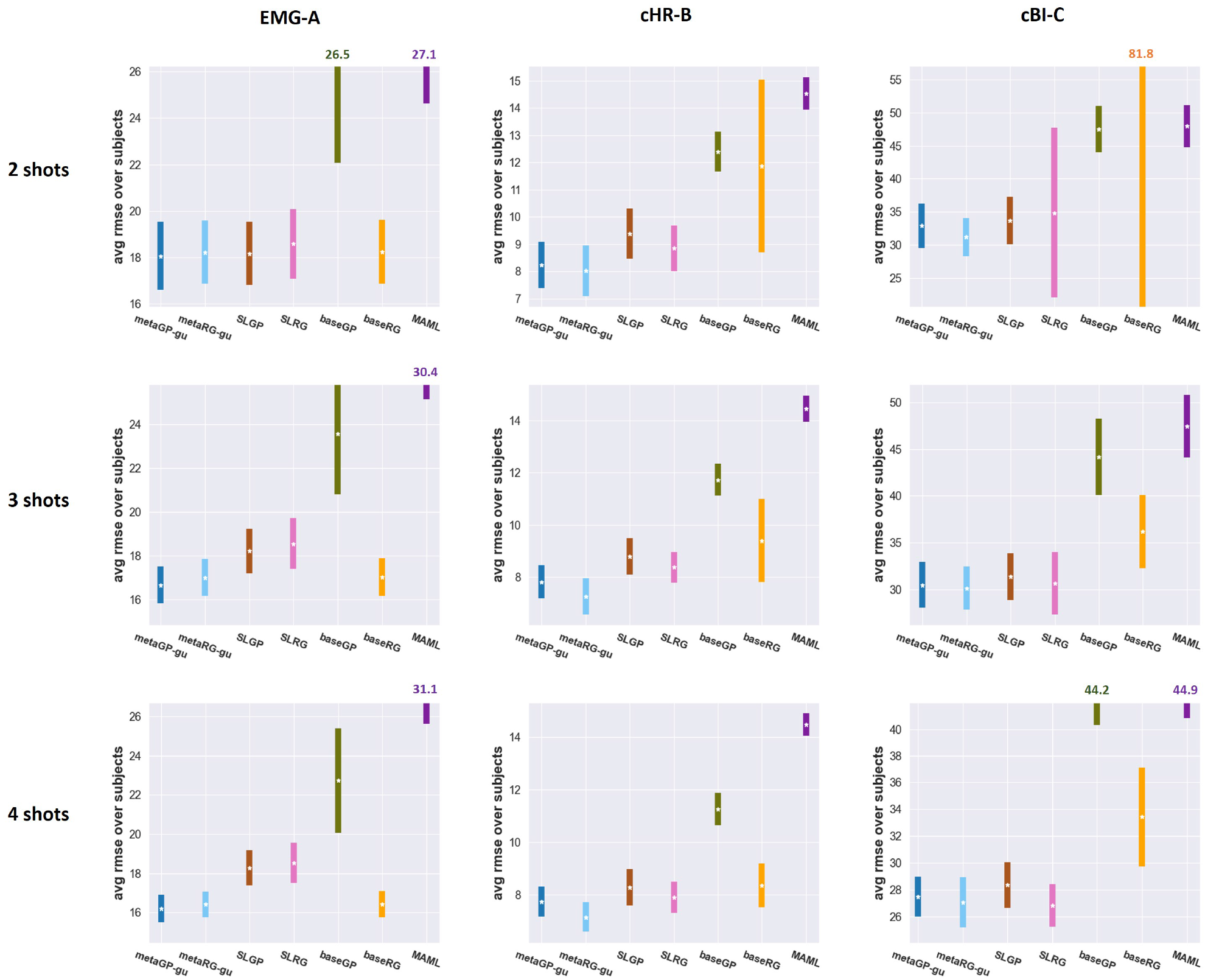
Performance comparisons on all three univariate datasets from Vagus nerve experimental data. For each dataset, leave-one-out cross-validation was conducted. The average meta-testing RMSE over all subjects are shown here as error bar plots, with their means represented as white star. For better visualizations, the error bars with large error are cut short, and its mean RMSEs are given on top of the error bars whenever the means are not shown in the plot. Please refer to Table B1, B2 and B3 for the complete quantitative results.

**Figure 5.**
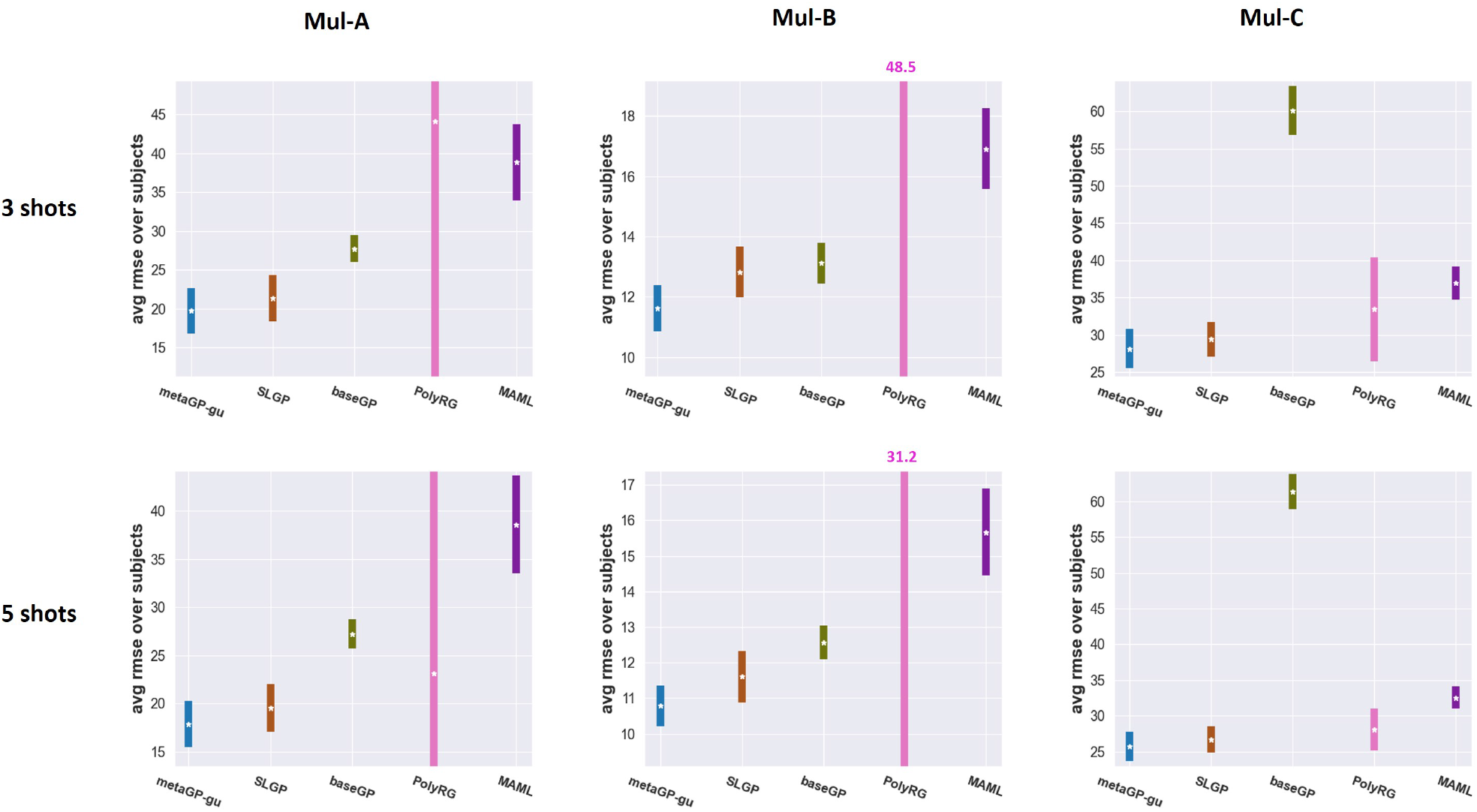
Performance comparisons on all three mulvariate datasets from Vagus nerve experimental data. For each dataset, leave-one-out cross-validation was conducted, and the average meta-testing RMSE over all subjects are shown here as error bar plots, with their means represented as white star. Same as Figure 4, the error bars with large error are cut short, with its mean RMSEs given on top of the error bars whenever the means are not shown in the plot. Please refer to Table B4 for the complete quantitative results.

Unlike the synthetic case, both `SLGP` and `SLRG` yield competitive performance in the real data, and `SLRG` slightly outperformed true meta-learning in one case (4-shot on `cBI-C`). Importantly, our framework has the better overall performance and is the most robust across all cases. Recall that the main difference between `SL-` and our meta-learning framework lies in the ways to infer the scaling factors: one with trivially using the maximum of few-shots to normalize data for a new subject, and the other by solving BO to infer this normalization. In some problem settings, naively normalizing by the max of a few-shots sample can be a good approach, but is far from being a principled approach. Conversely, our meta-learning framework consistently performs well across settings.

For better visualizing the derived individual fits, we provide in Figure 3 each subject’s averaged derived individual fits when given 3-shots, for both `metaGP-gu` and `metaRG-gu`. Judged by the curves, they are able to capture the varieties in the subjects’ operating ranges of both physiological and fiber responses’ amplitudes, and their derived individual fits correlated well with the data points.

Our proposed meta-learning framework also stands out for the multivariate datasets as consistently the best-performing one in all cases, and `SLGP` comes the second. Contrary to the performance of `SLRG` in the univariate cases, `PolyRG` exhibits a huge degradation and in particular suffers from a high variance in the performance. It is potentially due to the poor estimates from only the few-shots of data, especially on the quadratic term for each input where the “errors” on these estimates are also quadratically enlarged. Moreover, unlike the univariate datasets where each subject has either an exponential or linear input-output relationship, some input dimensions of the multivariate subjects do not demonstrate such clear correlations with the output, for example, between ∆*HR* and A fiber responses. This “weaker correlations” between input and output could lead `PolyRG` to overfit to the spurious relationship in the training data. On the other hand, the fact that `metaGP-gu` still performs well here further verifies its capability and flexibility to practical multivariate cases. We believe that reasons our meta-learning methods handled the weaker correlation better can be attributed to the use of GP, which innately has a consideration for uncertainties on the fits. However, we do need to point out that in the original paper reporting this VNS data [21], the coefficients of polynomial regression models were trained and further pruned based on statistical significance. Here we skipped that in our implementation of `PolyRG`, to keep consistent with the other methods in comparisons, as a thorough hyperparameter tuning was not applied on GP neither.

#### 2.3.3 Evaluation on skewed few shot samples of vagus nerve fiber recruitment

We now outline the central advantage of our meta-learning approach: fit estimation on biased few-shot samples. In neurostimulation use cases, practitioners might need to sample few stimulation parameters conservatively. For example, for safety concerns, one might want to limit an initial range of stimulation amplitudes. This practice is detrimental to the naive rescaling used by `SLGP` where normalization is done with the maximal query. As such, a method that can use few shots from a non-representative range of stimulation parameters and extrapolate consistently is an important asset. In the experiments presented so far, few shots were always uniformally sampled from the true operating range of a new subject. Now, we compare the performance of both *naive* and *meta* methods when 3-shots are sampled evenly from a decreasing *X* percentage of new subject’s data range. This is done for each subject by finding the data points which are *X* percentage smallest in terms of the charges and sampling the few-shots strictly from those data points. Note that since data of `Mul-C` has larger range for fiber responses, when conducting reduced percentage evaluation on `Mul-C`, we increased the BO’s search area by an additional 100 in each dimension’s upper bound. This is important especially for smaller percentage, because then the few-shots are further away from the true maximums. Please refer to Appendix A for more details regarding the implementation of BO. As the results shown in Figure 6, decreasing the percentage leads to consistent drop of `SLGP`’s performance. On the other hand, `metaGP-gu` is more robust to smaller percentage, since the BO inference model of `metaGP-gu` can estimate appropriate scaling factors beyond the mere maximums of the few-shots. Contrary to the stable performance on `Mul-A` and `Mul-B`, `metaGP-gu` on `Mul-C` suffers visibly from a reduced intensity range, making it a case where the capability of sampling few-shots from the right range is crucial to the overall performance. We note that the gap between the methods for `Mul-C` in the 30% to 60% range is surprisingly small. This is a phenomenon specific to this dataset, because for some of the subjects of `Mul-C`, the data points with smaller charge had fiber responses close to 0. Having the few-shots with those fiber responses makes it very difficult to deduce the curve of the fits. Overall, it is clear that `metaGP-gu` has advantage over `SLGP` on both its robustness to the few-shots bias and its prediction accuracy.

**Figure 6.**
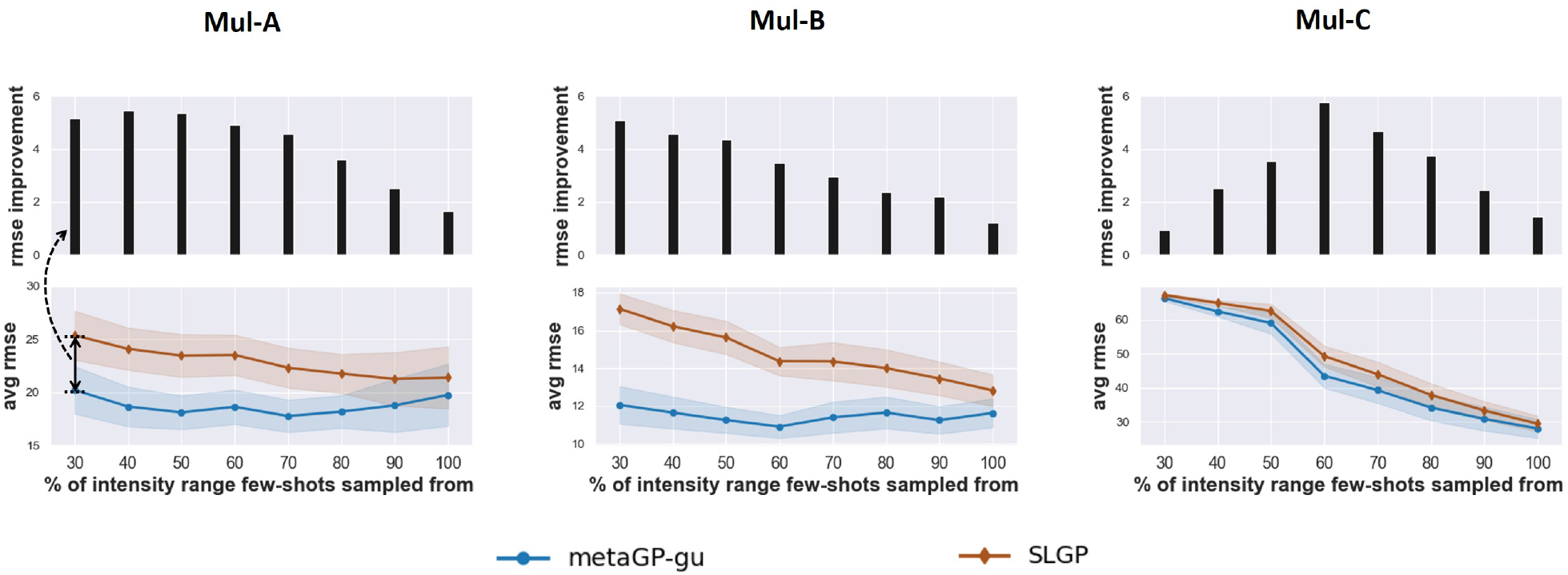
Performance comparisons between `SLGP` and `metaGP-gu` on all three mulvariate datasets from the real experimental data, in terms of different different percentage of the data range used in sampling few-shots. Note that for `Mul-C`, we increased the search range of BO accordingly in response to the larger range of its fiber responses.

#### 2.3.4 Evaluation of noise and data dimensionality with semi-synthetic datasets

We created a set of semi-synthetic datasets, hereon referred to in unity as `semi-C-syn`, as synthetic expansion of real dataset `cBI-C`. The procedure of generating semi-synthetic subject is similar to that of purely synthetic subject, except that certain statistics are derived from the real data. Please refer to Section 4.5 for details regarding the semi-synthetic datasets.

Using `semi-C-syn`, we probe how different aspects of the population sets could affect the few-shot prediction performance in `metaGP-gu`. We created multiple population sets varying four parameters defining the datasets, namely the number of subjects (`ns`), the number of data points from each subject (`np`), the noise level (`l_ni`) represented as a multiplier on the noise’ standard deviation when generating each subject, and the response range (i.e. fiber response) (`y_nm`) represented as a multiplier on the mean of distribution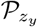 The population models trained on the various datasets are evaluated on the same meta-testing set to ensure fair comparisons. The results are shown in Figure 7. We can again observe the similar trends of better prediction when increasing the number of shots in the semi-synthetic datasets. However, despite being both synthetic datasets, unlike the situation for `syn-exp`, here the `metaGP-gu` approaches the dataset noise much slower and there remained sizable gap between the prediction error and the dataset noise. This is likely due to the discrepancies among the true generating functions of the synthetic subjects, which are sampled from GP population fits. In comparison, note that the subjects in `syn-exp` share exactly the same exponential function.

**Figure 7.**
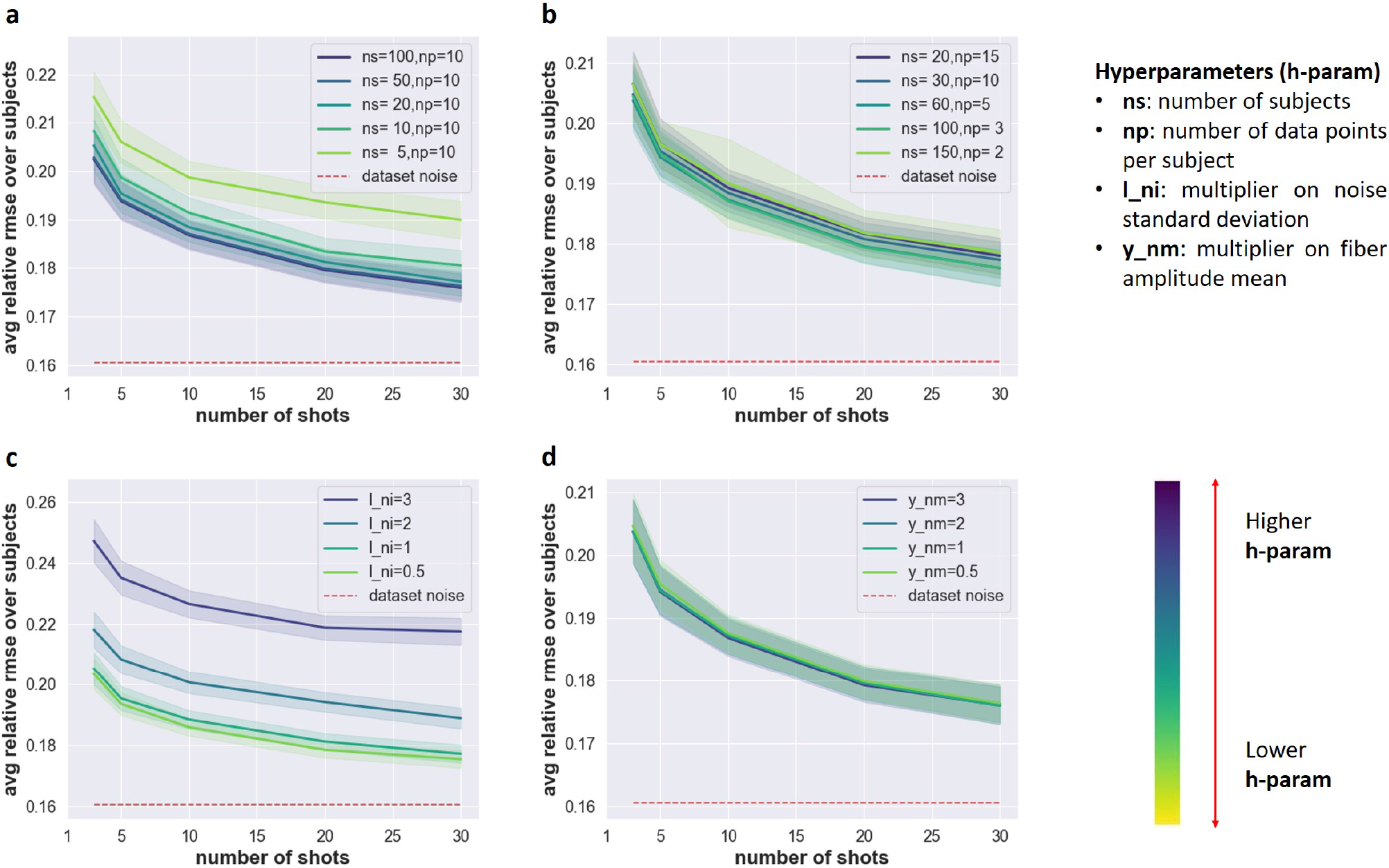
Investigations on the effect of population set on `metaGP-gu` fit with variations in four dataset parameters: (a) varying the number of subjects in the population set; (b) varying the number of data points in each each subject while keeping the total size of population set fixed; (c) varying the noise ratio of the population set; (d) varying the operating range of the population set.

Among the four dataset parameters, `ns` and `y_ni` prove to be important factors of the population set. This is reasonable as both the dataset parameters directly affect the population fit in terms of quantity and quality. In practice, noise level of the data is generally not directly manipulable, but this evaluation result suggests that increasing the data size can be an effective countering measure to obtain an informative population set. On the contrary, as shown in Figure 7(b) and (d), varying `np` and `y_nm` is not as influential. `np` directly affects the normalizing quality as we take the maximums in each subject as its normalization factors, so using a very small `np` would lead to poor estimates. In addition, `np` must not be 1 as it will make every data point normalized as 1. However, the results in Figure 7(b) also hints that the lack of data points in each subject could be complemented by having more subjects, as every dataset involved in Figure 7(b) had the same number of total data points (300). Lastly, the less important role of `y_nm` is easier to understand, as population fit is trained only by the normalized data, where the effect of a multiplied fiber amplitude was largely suppressed. Please see Appendix E for more discussions regarding `np` and `y nm`.

## 3 Discussion

The design principle of our proposed meta-learning framework is to transfer prior knowledge embodied in a neurostimulation population model into the learning process, so that the fast adaptation of an individual model is feasible given only very few data points from a new unseen subject. The advantages of our meta-learning framework are two-fold. First, using only few-shots, it is a data fitting tool for scientific investigations in experiments where varied subjects provide limited data. Second, it can provide information to help clinicians tune neurostimulation protocols in new patients by leveraging population data from past patients. These advantages are exemplified in the VNS fiber amplitude prediction problem, where we demonstrate our framework’s usefulness to investigate the relationship between fiber and physiological responses. Critically, for neurostimulation tuning, the ability of having less stimulations to achieve the same objective is generally preferred due to the safety concern for the subject. Hence it is beneficial to keep the repetitions of that process to a minimum. In addition, with the BO inference model, our method can work with the query data whose stimulation intensity is intentionally kept to a relative lower amount, a scenario that can occur in practice since the practitioner might want to act conservatively on the delivered stimulation intensity to the patients. Furthermore, our framework allows us to probe the trade-off between queries (shots) and performance. As shown in many evaluation results, more shots in general will lead to an improvement on the prediction error. Overall, this provides a tractable mechanism to adjust the performance based on both needs and constraints of practitioners: with more budget to collect data points, the methods would reward us with a better precision. Crucially, while there are situations where other fitting methods perform comparatively to our approach, we note that our meta-learning methods are consistently better than other methods overall, and that the advantage is increased in conservative procedures where safety concerns might lead to initially subsampling stimulation ranges for new subjects.

In the concrete problem of VNS, the core idea of the meta-learning framework is to leverage the shape of the population fit in the few-shot prediction of individual fiber recruitment by learning the subject-dependent operating ranges. This conveys a notion of knowledge transfer from the population to each new individual, which is specific to the problem but not restricted to the context, i.e. VNS. Since the challenges of personalization exist universally, our findings can be extended to other stimulation modalities and observables. In particular, our proposed meta-learning framework can be flexibly adapted to operate in many other problems similar to our case study, i.e. regression problems with subject-dependent scalings. One can simply replace the inputs and outputs accordingly across various settings. However, with subject-dependent parameters beyond scaling, modifications on the framework will be needed. In general, rescaling can be considered as a specific way to “condition” the population fit, so other approaches to provide conditioning can be explored. This entails also redefinition of the population model. Aside from being a population mean model, it can also be several layers of a deep learning model shared by all subjects, or a designated meta model solving a key sub or auxiliary problem. We leave such exploration to future works, but we will provide more discussions on modifying the framework in the next section.

The described meta-learning framework includes the option to utilize inner-loop feedback, in the form of one-step gradient update. In general, we can characterize the methods that use gradient update as the middle ground between placing full confidence in the prior knowledge from population and placing full confidence in the obtained few-shots from each individual. The former is closer to `SLGP` and `SLRG`, while the latter is functionally similar to `baseGP` and `baseRG` with a good initialization if we allow sufficient gradient steps until convergence. By performing the gradient update with an appropriate number of steps, the derived individual model has the potential to not only attend to the distinctness of each individual but also alleviate overfitting to the few-shots. Note that the latter is especially important for neural measurements, which commonly have large noise. In our implementation of the framework, we adopted one-step gradient update. Although the number of steps was not carefully tuned, the evaluation results justified the choice. However, the number of gradient update should be viewed as a domain-sensitive hyperparameter, and the exact number is subject to change for different problems.

A key advantage of the proposed meta-learning framework is its compatibility with GP as a generic task-agnostic population model for any future data. Whenever a good regression model is easily obtainable, like a close-form one, it could be ideal to use it. For example in the case of `syn-exp`, the exponential function is indeed the true data generating function hence the optimal choice for population model. But it is impractical to always assume having access to such a good regression model, and even there is, the required time dedicated to find it needs to be taken into considerations. In comparison, GP is advantageous in terms of flexibility, since it is a non-parameteric method whose fit depends on the data. GP also measures the uncertainties on the predictions, which has a good synergy when dealing with noisy neural measurements. We also note that the choice of population model goes beyond just GP and regression, in fact, any model with a tangible fit is directly applicable.

### 3.1 Limitation and Future Work

In its current form, our proposed meta-learning framework works the best for the task whose data is in general unimodal, since we assume that the shape of the learned population model is the extracted prior knowledge from the task. This can impede the future applicability of the framework, when the task is highly bimodel or multimodal, for example a mix of cubic and sine functions, in which case learning a single population model could be not instructive anymore, as it might eventually capture only one of the many modes. Handling multimodal task is also a problem of handling uncertainties and ambiguities in the few-shot learning, as one set of few-shots can belong to multiple models. Modifications of the framework to accommodate bimodel and multimodel in the population models are one important future direction.

Up to now we have been focusing on the correlation across different subjects in one task, and another direction to pursue is on the correlation across different tasks for one subject. Correlation among different fiber groups of the subject was possible in the currently used real datasets, because the amplitudes of A-, B- and C-fibers were all measured from eCAPs [21]. Although it remains unclear if learning on one fiber group would affect the others, multitask learning [30] can still be leveraged for such attempts.

In the last section, we mention the need to modify the framework to be more general, and that involves re-defining the key concepts in the framework. For example, to handle more diverse data formats, we can use convolutional networks handling the image inputs, recurrent ones for time-series signals or even a hybrid if the dataset is highly heterogeneous. Then one choice for subject-dependent parameters can be the coefficient and bias term for a linear modulation on the weight matrices, like the meta operation scaling and shifting proposed in Meta-transfer learning [26] and the feature-wise linear modulation layer from [27]. Powered by the use of deep learning, in this way the framework has the potential to be even more problem agnostic, but likely at the cost of less interpretable meta models and more data-hunger. Similar idea has been investigated in meta-learning literatures [29], but it remains an open question how to incorporate it for neurostimulation problems. Meta model can also be one targeting a different auxiliary problem shared by all subjects, where solving it can greatly aid in solving the main problem. However, the existence of such auxiliary problems is domain-specific, which needs to be investigated on a case-by-case basis.

Beyond regression problems, we can investigate the use of the meta-learning framework on other problem types, such as the searching problem for optimal stimulation parameters using BO as well as the sequential decision problem for closed-loop systems using optimal control and reinforcement learning [31, 32].

## 4 Methods

### 4.1 Problem Formulation for Few-shot Regression

In a typical supervised learning [33] setup of machine learning, we learn a model on a training dataset *D*_*train*_ and evaluate its generalization performance on a testing dataset *D*_*test*_. On the contrary, in meta-learning [10], we consider two meta datasets: meta-training set 𝒟_*meta_train*_ and meta-testing set 𝒟_*meta_test*_, each of which includes multiple regular datasets with its own *D*_*train*_ and *D*_*test*_.

We can formulate the few-shot prediction for individual subject’s fiber recruitment as k-shot regression, where the data points from the same subject construe a regular dataset *D* ∈ 𝒟 For each subject, k physio-fiber response pairs are made available to the model as *D*_*train*_, while the rest are used for evaluation as *D*_*test*_. For the typical few-shot learning setting, the value of k is small, for example *k <*= 5, while the larger value of k can be used to investigate the asymptotic performance. We consider the split of meta datasets by subject indices, meaning any subject is in either 𝒟_*meta_train*_ or 𝒟_*meta_test*_ but not both. Note that this is to minimize the information leakage from meta-training to metatesting.

### 4.2 Related Work on Personalized Treatment and Meta-Learning

#### 4.2.1 Personalized treatment

Instead of providing the same treatment for all the patients, personalized medicine aims to make appropriate adjustments for each individual, and it is a widely encountered and investigated topic in neurostimulation [1, 42–46]. For example in [1], the treatment devices for SCS were reprogrammed to tailor to each patient’s preference, and in [43], personalized information was incorporated into the BO algorithm as an extra input dimension for brain stimulation. The importance of personalized treatment is also highlighted in VNS, as the desirable stimulation parameters can be challenging to find due to subject variability in response to stimuli and various physiological factors [7, 18, 21, 23].

#### 4.2.2 Meta-learning

In the recent years, meta-learning for few-shot learning was widely studied on applications including but not limiting to computer vision [10], natural language processing [47], robotics [48] and drug discovery [49]. One taxonomy approximately categorize meta-learning algorithms into three classes: metric-based, optimization-based and model-based [50]. In general, metric-based algorithms map the input to a common feature space and make the prediction in the sense of weighted average. The weights correlate with the metrics calculated between pairs of inputs, measuring the similarities between them. For example for Matching Network [51], softmax of the cosine similarity between the embeddings of two inputs is used as weight on the predictions. On the other hand, optimization-based algorithms aims to learn as similar as an optimization process, and the challenges are to answer how the gradient calculated on the few-shots can lead to a good solution in a few optimization steps. An example of optimization-based algorithm is MAML, which tries to learn from meta-training set a good initialization on the model parameters, so that as small as one gradient update can let it adapt to a new task. Lastly for model-based algorithms, specially designed model are usually utilized for learning. In this category, MANN (Memory-Augmented Neural Network) [52] utilizes explicit external memory module to memorize the information and to facilitate fast learning of a new task. There is also natural connection between meta-learning and hierarchical Bayesian modeling, which utilizes a hierarchical form to infer the latent variables via Bayesian method [53], if viewing the meta-parameters and the model parameters as different level in the hierarchy. For example, [54] identified MAML can be seen as a point estimate version of empirical Bayes, a hierarchical Bayesian model whose parameters’ prior distribution is estimated using the data, and in [55], the authors showed the derived PAC-bayes bounds for meta-learning was similar in the expression as the one for hierarchical variational Bayes.

Note that our proposed meta-learning framework is closer to the optimization-based algorithms, where we extracted the learned population model as an “initial point” for each individual model and incorporated gradient graduate from both inner-loop and outer-loop error feedback. To the best of the authors’ knowledge, there hasn’t been work investigating the usage of few-shot learning and meta-learning for VNS and neurostimulation applications.

### 4.3 Gaussian Process

GP can be thought as a generalization of Gaussian probability distribution that describes functions instead of scalars or vectors [24]. GP is a Bayesian method, and it defines a posterior distribution over functions given the prior as well as the data. The prior represents the beliefs we have over the kind of function we are looking for, before even seeing any data. The belief will be updated and refined after observing data points, which leads to the posterior distribution.

For example, the posterior can put more preference on functions that better explains the observations. In addition, GP is a non-parametric model [9], whose capacity to fit increases along with more data points. When working with GP, we generally do not need to worry about underfitting the data, thus it is very flexible to use on any problem.

We consider GP for regression in our problem. Take a zero mean GP in the following form

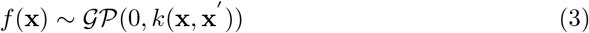

where *k*() defines the kernel function, which is the covariance function between two inputs **x** and **x***′* representing the covariance between their outputs. Therefore, given multiple training inputs **X** and its observed outputs **y**, assuming zero mean additive noise *ϵ ∼* 𝒩 (0, *σ*^2^), the predictive mean and covariance of testing inputs **X**^*∗*^ can be written as

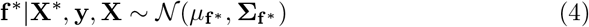

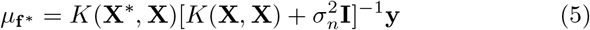

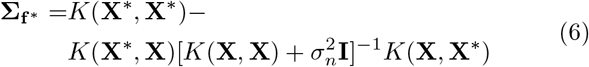

where *K*() is the covariance matrix between two sets of points. Note that both Equation (5) and Equation (6) contains matrix inverse operation which has *O*(*N* ^3^) complexity [34], where *N* is the number of entries in each dimension of matrix. As a result, GP is in general effective in small data domain but suffers from computational issue when a large data size is involved.

### 4.4 Bayesian Optimization

BO [35] is a method to find the maximum of a target cost function, especially practical when the one in question is expensive to evaluate. It does not rely on gradient, hence it is very useful when we do not have access to that information of the cost function. BO estimates the unknown cost function with a surrogate function, and like GP, it obtains posterior by updating the prior beliefs based on evidence, which is the responses of the function on multiple query points. Being both Bayesian methods, BO and GP share obvious synergy, and in fact, GP is a common choice of BO’s surrogate function. BO is an iterative process. At each iteration, after deriving the new posterior, BO utilizes an acquisition function to determine where to query the target function at the next step. The role of an acquisition function is to balance exploration and exploitation aspects when searching for the next query point. The exploration is to query the high uncertainty region, where we haven’t thoroughly investigated yet, and the exploitation is to query the high reward region, where we already observed higher values of responses from the function. By designing acquisition function to strike a profitable trade-off between exploration and exploitation, BO can keep the necessary queries to the function to a small number, which is essential when evaluating the target function is expensive.

Our proposed meta-learning framework applies BO to infer the scaling factors for each subject on both physiological and fiber responses’ amplitudes, based on its given k shots. Take 1-d regression problem as an example, we designed a scalar cost function for each candidate pair of (*z*_*x*_, *z*_*y*_). The cost function evaluated a scaling factors pair, by measuring the prediction error of the derived individual fit, obtained by rescaling the population fit with the said pair. We can write the corresponding optimization problem as:

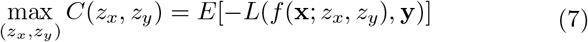

where *f* (**x**; *z*_*x*_, *z*_*y*_) = *z*_*y*_*f*_*ppl*_(**x**/*z*_*x*_), *f*_*ppl*_() and *f* () represent population model and derived individual model, and the cost function *L*() can take the form of RMSE and mean negative log-likelihood.

We adopted an improvement-based acquisition function, Expected Improvement (EI) [35, 36], to guide the search for the next query. EI works by finding the new query point by maximizing the expected improvement function

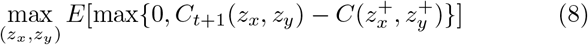

where 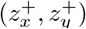 is the best scaling factors pair found so far. Furthermore, EI in Equation (8) can be evaluated analytically assuming probability density function and cumulative density function of the standard normal distribution [36].

### 4.5 Synthetic Data Generation of Purely Synthetic and Semi-Synthetic datasets

Algorithm 2 describes the synthetic subject generation procedure for both synthetic subjects. Note that although there are no specific meanings assigned to the inputs and the outputs of the synthetic datasets, we intend for them to mimic physiological and neural responses respectively.

#### 4.5.1 Purely Synthetic Datasets

Each subject in the purely synthetic dataset was generated by the following exponential function, which we will refer to as `exp`.

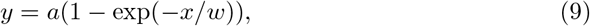

where *a* and *w* are the function parameters, corresponding to *z*_*y*_ and *z*_*x*_ in Algorithms 2 respectively. By varying values of *a* and *w*, the generation routine created synthetic subjects with different scalings on both input and output axes.

The `syn-exp` purely synthetic dataset contains 20 synthetic subjects generated from `exp`, where each subject has 200 data points. During subject generation, We restrained the domain of `exp` within [0, *w*]. However, note that due to additive Gaussian noise, it is possible that *y > a* and *y <* 0. Whenever k-shots are needed, a specific data space is divided into k mutually exclusive sub-region and one shot is sampled from each sub-region. For `syn-exp`, the data space is the input domain. Note that we introduce this practice so that the sampled few-shots are more evenly distributed and not clustered.

##### Algorithm 2 Data Generation Routine for Synthetic Subject i

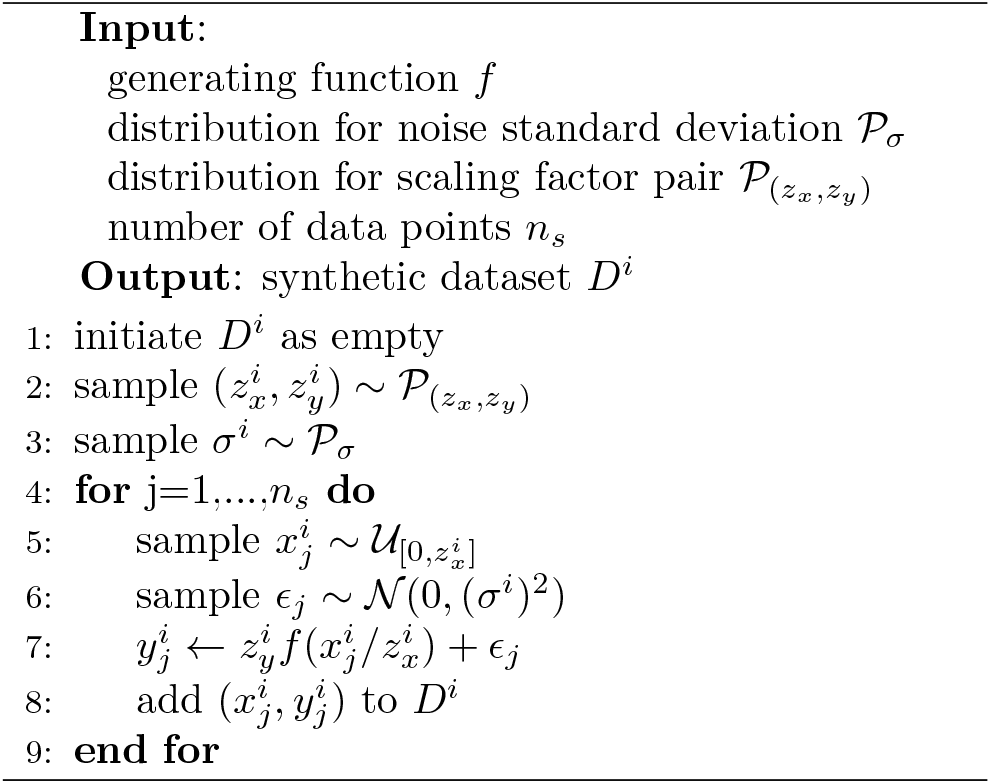

#### 4.5.2 Semi-Synthetic Datasets

As synthetic expansion of real dataset `cBI-C`, we used the statistics calculated from the real data to generate semi-synthetic subjects, so that the generated semi-synthetic subjects are reasonable to be appeared in practice:

- The generating function for each subject *f* ^*i*^ was sampled from population GP fit for `cBI-C`.
- Scaling factor 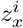 was sampled from Gaussian distribution, with 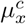 and 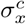 being the empirical mean and standard deviation of maximum physiological responses across real subjects. The same strategy was repeated for 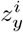 on fiber responses.

Noise standard derivation for each subject *i* was computed as 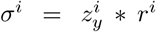 where the process to obtain *r*^*i*^ was described as follows. Firstly, individual model was fitted for each real subject *m*. In other words, we fitted for each subject with `exp` using all its data points. Secondly, we recorded the ratio of the training error to the maximal range for each real subject *m*, written as 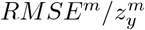 as an approximate noise-to-signal ratio of that subject. Finally, *r*^*i*^ was sampled from another Gaussian distribution, with *µ*_*r*_ and *σ*_*r*_ being empirical mean and standard deviation of the above-mentioned ratio across real subjects.

To bear resemblance to practical situations, we kept the number of data points per subject relatively small when creating the set of semi-synthetic datasets `semi-C-syn`, whilst including a larger number of subjects. Sampling scheme of semi-synthetic datasets for few-shots is same as that of purely synthetic datasets, and we report the few-shot prediction performance of the same meta-testing set while varying the population sets.

Following the above-described procedure, the created meta-testing set contains 50 synthetic subjects and 50 data points per subject. In addition, we constructed a set of semi-synthetic datasets sololy as the population sets, with variations in the following four dataset parameters: number of subjects (`ns`), number of data points per subjects (`np`), responses’ noise level (`l_ni`) and range of fiber amplitude (`y_nm`). Among the four, the former two control the population set in quantity, while the latter two correspond more to the variations in quality. In particular, to test on `ns`, we created a semi-synthetic population set of 110 subjects, with 10 data points per each subject. Then for different values of `ns`, we sampled from this population set different numbers of subjects, ranging from 5 to 100. For each value of `l_ni`, we recreated a similar population set as the `ns` one, but replacing noise standard deviation *σ* with *λ*(*z*_*y∗r*_), with *λ* as the additional multipliers ranging from 0.5 to 3. When testing those population sets for `l_ni`, we sampled 20 subjects from each generated population set. The similar procedure was repeated for `np` and `l_nm`, where their population set has 160 subjects, each with 20 data points. To test on `np`, we included different combinations of subject number and points number per subject with `np` ranging from 2 to 15, while maintaining the total data points fixed as 300. As for `y_nm`, we created different population sets as the one used for `np`, while varying the multipliers on ranges of Y-axis (fiber response) from 0.5 to 5, and we sampled 20 subjects and 15 points per subjects from each generated population set.

### 4.6 Brief Overview of Real Experimental Data

The utilized real experimental datasets are from publicly available data introduced in [21], including one set with univariate inputs and the other with multivariate inputs. In both cases, the data was from rat experiments, and eCAPs were recorded while registering three acute physiological markers to the VNS. To keep the manuscript concise, here only the aspects of the real data pertaining to our evaluations are included, and for comprehensive descriptions regarding the data collection processes and further data analysis, please refer to the original paper.

The main challenges of few-shot prediction on the experimental datasets are subject-dependent scalings and the limitation on training size. For the former, the large difference on the personalized scaling difference deteriorates the heterogeneity within the datasets and thus must be attended to. Moreover, many existing meta-learning methods don’t focus on this aspect. The latter is likely due to the combinations of the difficulty of conducting the experiments, the maximum number of simulations can be delivered safely and the need to filter out invalid data points. The limited size of training data hinders the use of powerful tools such as deep learning, due to the risk of overfitting.

#### 4.6.1 univariate datasets

There are three univariate datasets `EMG-A` (relating EMG and A-fiber), `cHR-B` (relating changes in HR and B-fiber) and `cBI-C` (relating changes in BI and C-fiber), and the authors of the original paper identified a linear relationship for `EMG-A` and exponential ones for `cHR-B` and `cBI-C`. All three datasets are coming from 11 rat subjects, and the average data points per subject are 32.8 (ranging from 14 to 66) for `EMG-A`, 27 (9 to 54) for `cHR-B` and 12.8 (7 to 20) for `cBI-C`. Each of the datasets includes data from 8 subjects. The original paper described a closed-form exponential regression model in the form of `exp` in Equation (9), for the pair of changes in HR and B-fiber and the pair of changes in BI and C-fiber, and a closed-form linear model *y* = *Ax* for the pair of EMG and A-fiber, where *x* and *y* are normalized values over each subject’ respective maximal values.

Following the data preprocessing procedure in [21], We extracted the same datasets for our evaluations. The same closed-form functions are used as the regression models, and the same normalization scheme is adopted whenever needed. The utilized few-shot sampling procedure is the same as the synthetic cases but with feature charge-per-pulse, calculated by multiplying stimulus intensity by the pulse width. Since charge-per-pulse is purely defined by the stimulation parameters, it is a controllable feature by the practitioners. Therefore, it is reasonable to assume that we can query the few-shots based on the change-per-pulse feature.

#### 4.6.2 multivariate datasets

There are three multi-variate datasets which contain the data mapping from all three above-mentioned physiological markers and *charge-per-pulse* to each fiber group. Among them, `Mul-A` contains the data from 9 subjects with in average 29.1 points per subject (ranging from 14 to 54), `Mul-B` contains 8 subject with 30.6 per subject (18 to 53), and `Mul-C` contains 8 subjects with 31.5 per subject (17 to 54). The original paper described the training of a second-order polynomial regression model with these three datasets:

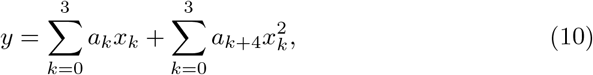

where *y* is the normalized fiber responses’ amplitudes, *x*_0_ is the charge-per-pulse, *x*_1_ is the normalized EMG, *x*_2_ is the ∆*HR*, and *x*_3_ is the ∆*BI*.

We used the same data points in our evaluations, but with zero entries replaced by 1*e*^−5^ due to the numerical issues of rescaled by 0. Among the methods that work with normalized data, we applied normalization to all dimensions of the multivariate inputs, for `metaGP-gu` and `SLGP`, and to only EMG for the polynomial regression `PolyRG`. For the three physiological markers and the fiber responses, the normalization was conducted in the same way as the univariate cases, with maximal values within each subject, but for the charge-per-pulse, we used a constant value 6000. Lastly, for `MAML` we normalized the charge-per-pulse to stabilize the learning process.

### 4.7 Implementations

We implemented the proposed meta-learning framework in Python. PyTorch [37] packages BoTorch [38] and GpyTorch [39] were utilized to perform BO and GP respectively, whereas all kinds of closed-form regressions were optimized with `scipy.optimize.curve fit` [41]. We used `scipy.stats.gamma` as gamma pdf when generating the multivariate synthetic datasets, and `Torchmeta` [40] library for `MAML`. We adopted PyTorch’s automatic differentiation engine `torch.autograd` for all gradient computations. All the evaluations were ran in CPU machine, except for `MAML`, which were ran in a single GPU with 48G memory.

## Acknowledgement

This work is supported by grants to GL from Canada CIFAR AI Chair program, the Canada Research Chair in Neural Computations and Interfacing, and the Fonds de Recherche du Québec (research scholar program), and a grant to SZ from United Therapeutics Corporation (MD, US). The authors would like to thank Theodoros Zanos, Numa Dancause, Marco Bonizzato, Tristen Deleu and Leo Choiniere for discussions on the project motivations and future directions, Lorenz Wernisch for suggestions on related field, and Eric Elmoznino and Alexandre Payeur for suggestions on manuscript structures.

## Appendix A. Additional details on implementations

Here we provide additional implementations details for our proposed meta-learning framework and `MAML`. For the BO inference model of the meta-learning framework, we extracted the maximum of the few-shots and set the search range for each individual dimension to be [0.5 *∗max, max* + 50]. The budget of points to search in BO is 25, including 10 random initial points and 15 others in the process, and we skipped the new points to search when it was too close to the existing ones. In practice, the BO process can fail due to numerical issues, especially with a higher dimensional search space, so we allowed the process to repeat up to 20 times if such error was thrown. The gradient update was performed via stochastic gradient descent (SGD) with a predetermined fixed learning rate, which we refer to as the inner-loop learning rate (ILR). We treated ILR as hyperparameter in our implementation and tuned it by running the few-shot prediction on the training data. The utilized value of ILR is dataset-dependent in range [0.05, 0.5].

For the implementation of `MAML`, we used multi-layer preceptron (MLP) with 2 hidden layers of size 40. The meta optimizer is Adam [56] with 1*e*^−3^ learning rate, and the inner loop optimizer is SGD with 1*e*^−5^ learning rate. We set `MAML` to use only the first-order gradient, as we found that using the Hessian was costly and didn’t change much of the performance. We found that `MAML` can diverge in training especially on higher multivariate datasets, so we introduced an early-stopping schedule to end the training loops when the mean training error of the past 5 epochs was larger than previous best plus 0.4. We found this scheduler capable of exit the training loop at reasonable time before severe divergence.

For all the reported performance in purely synthetic and real experimental data, we repeated 500 times per subject for `MAML` and 200 times for all others. The higher number of repetitions for `MAML` was because it tended to have unstable results. For the semi-synthetic data, we repeated 400 times per subject due to the sampling on the subjects and data points in each subject. With the per-subject mean and standard deviations, we compiled and reported the global statistics with the mean of all per-subject means and the square root of all per-subject standard deviation means 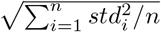 where *std*_*i*_ represents the standard deviation of subject *i*.

## Appendix B. Full evaluation table on real experimental univariate datasets

The full evaluation tables for real univariate datasets `EMG-A`, `cHR-B`, `cBI-C` and multivariate datasets `Mul-A|B|C` are shown in Table B1, Table B2, Table B3 and Table B4, respectively.

## Appendix C. Evaluations on multivariate synthetic datasets

### Appendix C.1. Multivariate Synthetic Datasets

For a synthetic subject with multivariate input, we consider the case where the product of independent components at each input dimension yields the output:

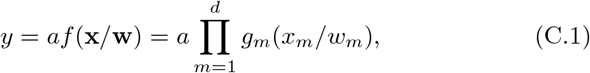

where *y∈* ℛ **x** *∈* ℛ^*d*^ and *g*_*m*_ ℛ*→* ℛ represents a 1-d function for *m*-th dimension of input **x**. As with other synthetic datasets, we set the domain of each *g*_*m*_ as [0, 1], then **w** *∈* ℛ^*n*^ are the scaling vector of the input, a generalization of *z*_*x*_ (in Algorithm 2) to multi-dimensional space. We consider two choices for *g*(·), one is the logistic function and the other probabilistic density function of gamma distribution, referred to as `logi` and `gam` respectively:

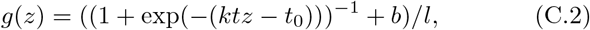

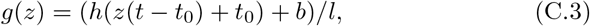

where *h*(*z*) = *z*^*α*−1^ exp^−*z*^/Γ(*α*), *t*_0_ controls the shifting, *l* is a constant to ensure *g*(*z*) is upper-bounded by 1 for *z* ∈ [0, 1], *b* is the bias, and *k* and *α* are function parameters corresponding to slope and shape respectively. Lastly, *t* is a specific one used to tune the effective domain of the 1-d function for reasons described below. Note that, jointly with *t* and *t*_0_, we can effectively “zoom-in” or “zoom-out” the curve of the 1-d function.

Due to the use of product in Equation (C.1), one challenge with this formulation is that for a higher dimensional input space, its output could be exponentially larger or smaller than its lower dimensional counterpart, i.e. either exploding or vanishing. Both exploding and vanishing can co-exist in a specific subject, creating a delta function like output structure, specifically in higher dimensional dataset. It is for this reason we introduce the normalization factor *l* to properly upper-bound each *g*(*z*) by 1, effectively decoupling the output scaling *a* from the input dimension. However, this doesn’t solve the vanishing issue, as normalizing multiplicands results in the output closer to 0 as the input dimension increases. This is undesirable as it would make higher and lower dimensional datasets incomparable. Therefore, we would desire the capability to maintain the appropriate and stable variance within the unscaled outputs, i.e. *f* (·) in Equation (C.1), and for that we brought in *t* and *t*_0_. Importantly, tuning *t* and *t*_0_ gains us the ability to effectively zoom-in specific region in the multivariate function’s range, so that we can stabilize the output variance across different input dimensions.

**Table B1:**
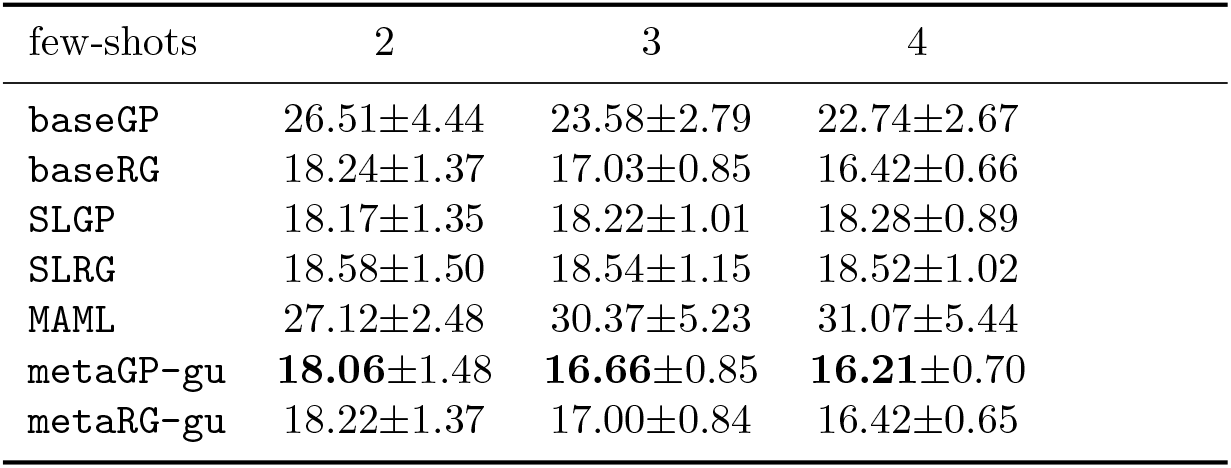
Full performance comparisons on EMG-A with average meta-testing RMSE and standard error over all subjects. The best performance is made bold in the table.

| few-shots | 2 | 3 | 4 |
| --- | --- | --- | --- |
| baseGP | 26.51±4.44 | 23.58±2.79 | 22.74±2.67 |
| baseRG | 18.24±1.37 | 17.03±0.85 | 16.42±0.66 |
| SLGP | 18.17±1.35 | 18.22±1.01 | 18.28±0.89 |
| SLRG | 18.58±1.50 | 18.54±1.15 | 18.52±1.02 |
| MAML | 27.12±2.48 | 30.37±5.23 | 31.07±5.44 |
| metaGP-gu | **18.06±1.48** | **16.66±0.85** | **16.21±0.70** |
| metaRG-gu | 18.22±1.37 | 17.00±0.84 | 16.42±0.65 |

**Table B2:**
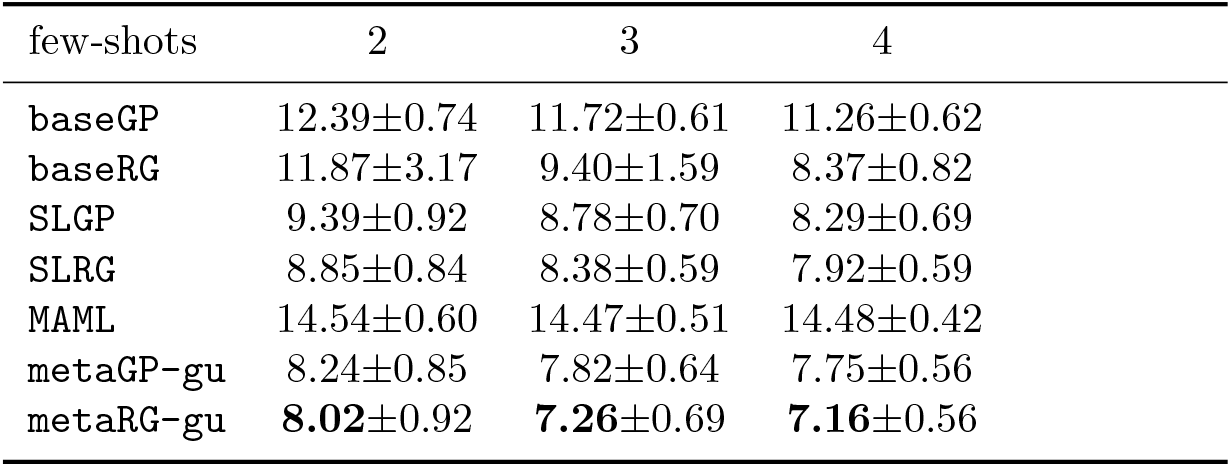
Full performance comparisons on cHR-B with average meta-testing RMSE and standard error over all subjects. The best performance is made bold in the table.

| few-shots | 2 | 3 | 4 |
| --- | --- | --- | --- |
| baseGP | 12.39±0.74 | 11.72±0.61 | 11.26±0.62 |
| baseRG | 11.87±3.17 | 9.40±1.59 | 8.37±0.82 |
| SLGP | 9.39±0.92 | 8.78±0.70 | 8.29±0.69 |
| SLRG | 8.85±0.84 | 8.38±0.59 | 7.92±0.59 |
| MAML | 14.54±0.60 | 14.47±0.51 | 14.48±0.42 |
| metaGP-gu | 8.24±0.85 | 7.82±0.64 | 7.75±0.56 |
| metaRG-gu | **8.02±0.92** | **7.26±0.69** | **7.16±0.56** |

**Table B3:** Full performance comparisons on cBI-C with average meta-testing RMSE and standard error over all subjects. The best performance is made bold in the table.

| few-shots | 2 | 3 | 4 |
| --- | --- | --- | --- |
| baseGP | 47.53±3.49 | 44.19±4.07 | 44.20±3.88 |
| baseRG | 81.81±99.56 | 36.19±3.92 | 33.44±3.72 |
| SLGP | 33.69±3.60 | 31.39±2.52 | 28.37±1.71 |
| SLRG | 34.85±12.82 | 30.68±3.31 | **26.85±1.59** |
| MAML | 47.97±3.15 | 47.42±3.33 | 44.89±4.07 |
| metaGP-gu | 32.92±3.33 | 30.49±2.43 | 27.48±1.50 |
| metaRG-gu | **31.20±2.85** | **30.16±2.31** | 27.07±1.84 |

**Table B4:** Performance comparisons on all three multivariate datasets from the real experimental data. For each method and each dataset, leave-one-out cross-validation was conducted, and the values reported here are its average meta-testing RMSE over all subjects. The best performance of each task is made bold in the table.

| few-shots | Mul-A |  | Mul-B |  | Mul-C |  |
| --- | --- | --- | --- | --- | --- | --- |
|  | 3 | 5 | 3 | 5 | 3 | 5 |
| baseGP | 27.75±1.70 | 27.23±1.53 | 13.12±0.67 | 12.56±0.48 | 60.12±3.31 | 61.38±2.52 |
| MAML | 38.85±4.88 | 38.59±5.07 | 16.91±1.34 | 15.67±1.22 | 36.95±2.19 | 32.55±1.56 |
| PolyRG | 44.10±87.69 | 23.09±37.46 | 48.49±52.54 | 31.19±39.35 | 33.46±6.97 | 28.05±2.89 |
| SLGP | 21.39±2.94 | 19.50±2.48 | 12.82±0.84 | 11.61±0.73 | 29.42±2.31 | 26.69±1.81 |
| metaGP-gu | **19.74±2.93** | **17.84±2.41** | **11.63±0.76** | **10.79±0.58** | **28.15±2.62** | **25.69±2.05** |

**Figure C1:**
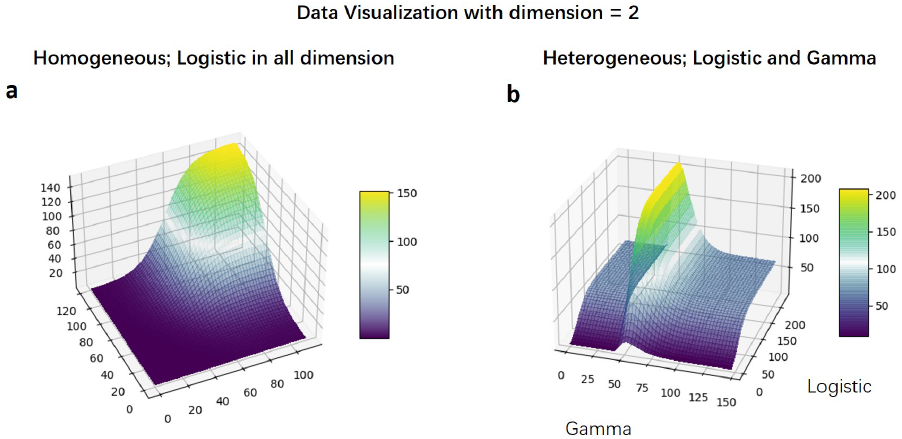
Illustrations on multivariate synthetic subject that is (a) homogeneous (b) heterogeneous.

Three multivariate synthetic datasets with multivariate inputs were generated for this evaluation. The first one is unimodal homogeneous, whose synthetic subjects all have the same homogeneous set of `logi`s as its *g*_*m*_*s*, and we refer to as `syn-muli-uniho`. Both the second and the third ones are heterogeneous, where for each of their input dimensions, it can be either `logi` or `gam` as *g*_*m*_. The second dataset is also unimodal, whose synthetic subjects share the same heterogeneous set of *g*_*m*_*s* (function type in each dimension and the function parameters), whereas the third one is multi-modal, where the set of *g*_*m*_ are sampled individually for each synthetic subject. Note that we refer to the second dataset as `syn-muli-unihe` and the third as `syn-muli-mulhe`.

The generation routine described in Algorithm 2 can be generalized to the multivariate case, by repeating row 5 on each of the input dimensions. As discussed above, when generating the multivariate datasets, we exhaustively searched for the combinations of function parameters, so that across different input dimensions both the variance of the normalized output and the mean of subject-to-subject unnormalized difference were kept to a similar level. The exhaustive search was conducted by iterating over a range of random generator seeds and sampling the function parameters from predetermined probability distributions. In addition, for the two heterogeneous datasets, they are designed that the number of `gam`s is always no less than and at most one more than that of `logi`s. For example, for dataset of dimension 5, there will be 3 `gam`s and 2 `logi`s, and for dataset of dimension 4, 2 for each. Figure C1 illustrate one of each synthetic subject from `syn-muli-uniho` and `syn-muli-unihe` with input dimension being 2.

Other than scaling performance to the multivariate input, we also investigated on the multivariate output. To decouple out the effect of multivariate input, we fixed input dimension as 1 and passed the inputs to three 1-d functions `exp`, `logi` and `gam` separately. The independent outputs from the functions constitute the multivariate outputs. We refer to this dataset as `syn-mulo`.

### Appendix C.2. On the scaling performances of multivariate input

Figure C2 shows the performance comparison on all three multivariate input synthetic datasets with 5, 10 and 20 shots from each new subject. Note that here we exclude evaluations below 5 shots as for even fewer shots, the risks of having all negative values for the outputs of the sampled shots is significantly higher, and this is especially a commonly-seen issue for datasets with higher dimensional input. The negativity on the outputs is a false reflection of the ground truth as all the underlying data-generating functions are with both positive domains and ranges, and the negative values exist only because of the added noise to the outputs. In general, we find the negativity issues rarely seen on 5 shots and above. Lastly, with input dimension being 1, the subjects in considerations are essentially univariate subject, similar to those in `syn-exp`. However it is expected to have discrepancies in their respective results since the subjects here have distinct data-generating function and larger scaling factors in general.

Judging from Figure C2, overall `metaGP-gu` demonstrates competitive performance when scaling to higher dimensional inputs. Despite a slightly higher error associated with the higher dimensional datasets, our proposed meta-learning method generally exhibit stable performance across different numbers of input dimensions. In general, among the three synthetic datasets, the best performance of `metaGP-gu` is found for unimodal homogeneous subjects, then unimodal heterogeneous ones and lastly multimodal heterogeneous ones. This is generally in line with the difficulties of the associated datasets, where the heterogeneity adds the non-monotonicity from the `gam` and the multimodality adds the subject variations. Furthermore, to better quantify the performance gaps among different scenarios, we show in Table C1 the average rate of improvement on rRMSE of `metaGP-gu` over other baselines for different number of few-shots. Compared to both `baseGP` and `MAML`, `metaGP-gu` exhibits clear advantages in terms of the prediction error, with a larger gap when more shots from the new subjects are accessible. Despite the overall poorer performance, `MAML` can still be competitive when the number of shots is small, for example in `syn-muli-mulhe` with 5 shots and input dimensions between 2 and 7. This is attributed to the few-shot prediction loops `MAML` runs with meta-training datasets which has the potential to extract prior knowledge more targeted to the overall few-shot learning procedure. Meanwhile, the results show that `MAML` doesn’t benefit as much as `metaGP-gu` from accessing more shots in the homogeneous datasets, and even suffers from having more shots in some cases of the heterogeneous datasets. Note that the degradation is likely caused by unstableness during training, which we will discuss in more details later.

**Figure C2:**
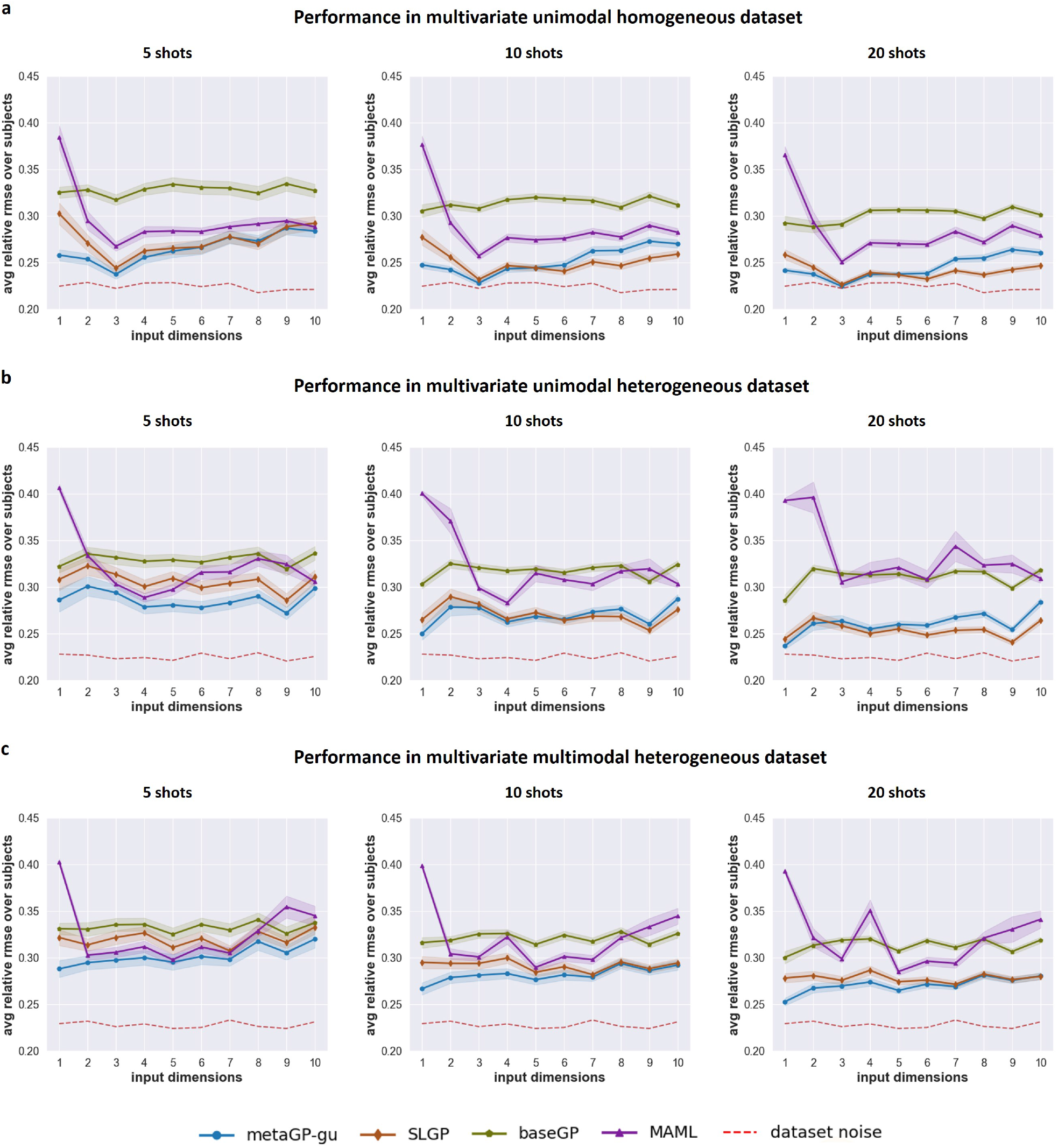
Performance comparisons on all three synthetic datasets with multivariate inputs. Plots show performance of all the methods in comparison on three multivariate synthetic datasets with input dimensions from 1 to 10 and with 5, 10 and 20 shots. (a) Performance on multivariate unimodal homogeneous dataset `syn-muli-uniho`; (b) Performance on multivariate unimodal heterogeneous dataset `syn-muli-unihe`; (c) Performance on multivariate multimodal heterogeneous dataset. `syn-muli-mulhe`.

Different from all other baselines, `SLGP` performs relatively better with more shots and with higher input dimensions. Recalls that `SLGP` rescales fitted population model naively with maximums of the few-shots. Given more shots from the subject therefore enables a more accurate estimate of the ground-truth values, and the problem becomes closer to the standard supervised learning problem with the maximum normalized datasets. Moreover for unimodal subjects, this exactly mimics how it is generated in the first place, so a superior performance of `SLGP` is expected with larger number of shots in both `syn-muli-uniho` and `syn-muli-unihe`. However, since that advantages rely on the accurate approximate of the true maximums, there will be negative impact when we couldn’t sample the full range. See Figure C3 where we conducted similar evaluations as with multivariate real data, on the two unimodal datasets, with decreasing percentage of the data range used in sampling few-shots. Due to a similar reason as `Mul-C`, we increased the BO search range by 150 in each dimension, in order to accommodate the significantly enlarged gap for smaller percentage, between the maximums of the sampled few-shots and the true maximums. We can observe the similar trend in Figure C3, where the relative performance of `metaGP-gu` to `SLGP` is better in smaller percentage. On the contrary for multimodal dataset where it cannot mimic the true data-generating functions, its advantages over `metaGP-gu` dwindle quickly even for higher dimensional datasets. Lastly, note that when the number of shots is limited, `metaGP-gu` still outperformed `SLGP` on all three multivariate synthetic datasets, justifying that correctly estimating the scaling factors is the bottleneck in this few-shot setting.

One interesting point observed from Figure C2, is that multiple methods display a non-monotonic behaviour with respect to the input dimension in the subfigures. In other words, the methods perform the best in datasets having middle value of input dimension. This is rather counter-intuitive since as a higher dimensional input usually means a larger hence more difficult space to navigate. We believe this phenomenon is related with how we generate these three multivariate datasets. As mentioned in the previous section, to keep fair comparison, we aim to keep a stable level of output variance across different input dimensions when generating the datasets. However, due to the product and the normalization, keeping the same output variance effectively means exponentially smaller output variance of each dimension’s 1-d function. In other words, if viewing from the perspective of a specific dimension, the 1-d function gets flatter and easier. The overall difficulty of the datasets associated to different input dimension is potentially a trade-off between these two factors: larger space to explore for higher input dimension and flatter function to have for higher input dimension, which could explain why the middle value becomes the best in certain cases. This characteristic of the synthetic datasets influences `SLGP` more than others, as it is easier to encounter good scaling factors estimate from few-shots when the overall function is flatter. This can also partially explain `SLGP`’s good performance in higher dimensional datasets.

Lastly, as mentioned above, `MAML` sometimes exhibits decrease in performance when given more shots. In our evaluations, this phenomenon occured usually for heterogeneous datasets and with higher dimensional input. Note that `MAML` operates on the unnormalized data as it doesn’t contain an explicit population model, hence there will generally be larger gradient propagated back to the model. When the model has access to more shots, it will have more information to exploit during the meta-training, and that, when combined with large gradient, could lead to more severe overfitting to the few-shots. In fact, more training divergence for `MAML` is in general observed on dataset of higher dimensions and more shots, and early stopping has to be applied during training. Although such undesirable phenomena could occur in both homogeneous and heterogeneous datasets, overfitting in the heterogeneous one could be more detrimental as it means to misplace the peak in `gam`, whereas the monotonicity and saturating aspects of `logi` in all input dimensions of `syn-muli-uniho` allows it to suffer less.

### Appendix C.3. On the scaling performances of multivariate output

Since each output dimension of subjects in `syn-mulo` are independently generated, we adopted the independence assumptions into the evaluations. In other words, for all the baselines, we treated each subject in `syn-mulo` as three independent 1-d univariate relationships sharing their input domains, and we reported the prediction performance as the average of the three. Note that this average value can also be viewed as the averaged performance of three different datasets with output dimension being 1 or even 2, as long as the proportion of subjects among the three functions is pre-served.

**Table C1:** Comparisons on the rate of average rRMSE improvement of our proposed meta-learning method over the baselines with 5, 10 and 20 shots, over all input dimensions.

| few-shots | syn-muli-uniho |  |  | syn-muli-unihe |  |  | syn-muli-mulhe |  |  |
| --- | --- | --- | --- | --- | --- | --- | --- | --- | --- |
|  | 5 | 10 | 20 | 5 | 10 | 20 | 5 | 10 | 20 |
| baseGP vs metaGP-gu | 0.191 | 0.197 | 0.184 | 0.132 | 0.150 | 0.159 | 0.093 | 0.122 | 0.136 |
| SLGP vs metaGP-gu | 0.031 | -0.007 | -0.019 | 0.065 | 0.001 | -0.030 | 0.057 | 0.033 | 0.027 |
| MAML vs metaGP-gu | 0.096 | 0.118 | 0.132 | 0.105 | 0.152 | 0.210 | 0.069 | 0.115 | 0.155 |

**Figure C3:**
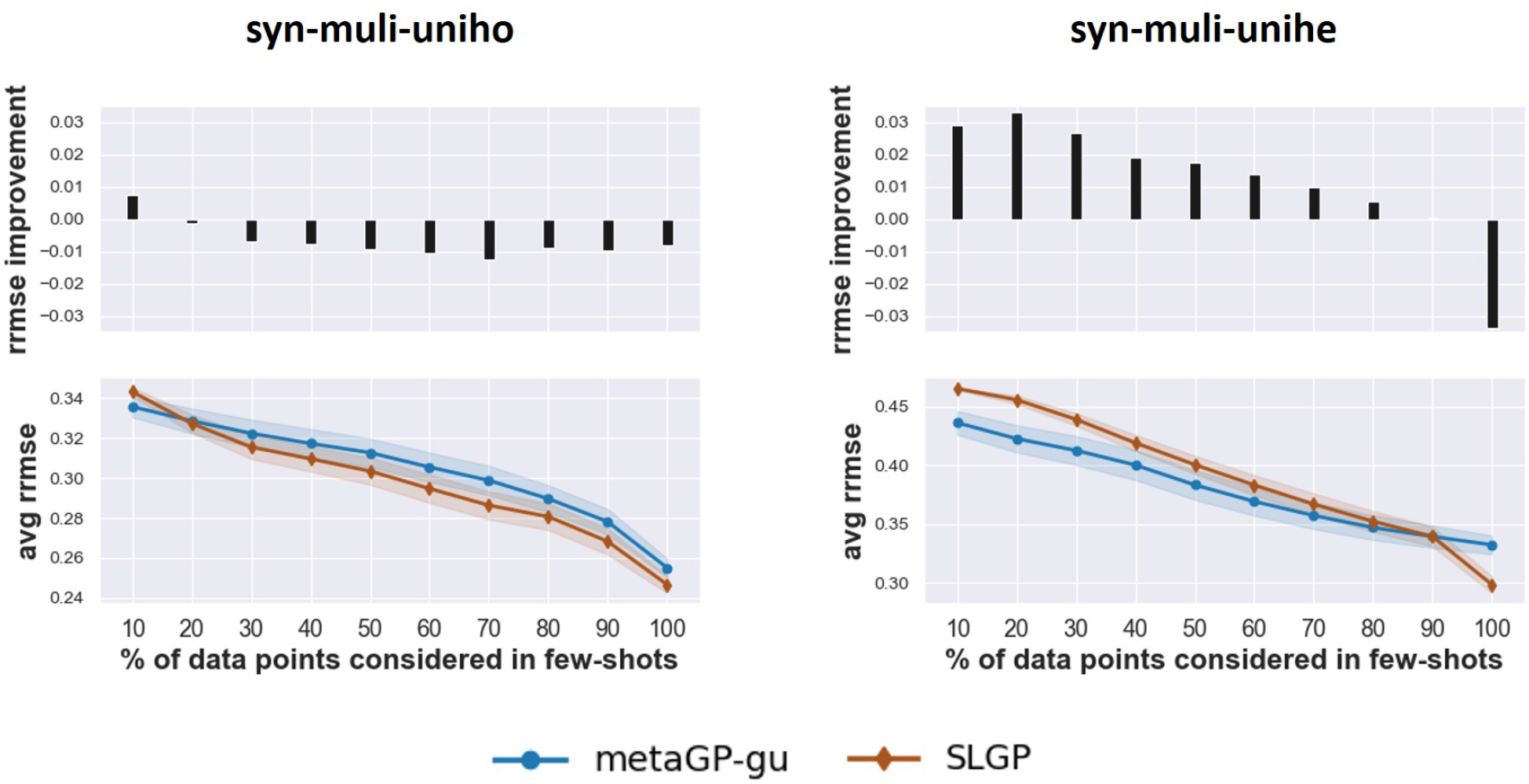
Performance comparisons between `SLGP` and `metaGP-gu` on both unimodal synthetic datasets with multivariate input, in terms of different different percentage of the data range used in sampling few-shots. Note that here for both of them, we increased the search range of BO accordingly in response to the larger range of its outputs.

However, the situation is more complicated for our meta-learning framework when inferring the scaling factors for each of the output dimensions. Consider one subject with the mapping ℛ → ℛ^3^, we would need one scaling factor for each of the input and output dimensions. However, if we treat this subject as three independent 1-d univariate relationships, there could be at most 3 scaling factors for the input (one for each output dimension), which contradicts with the fact that there should be only an unique input scaling factor. Hence for each subject with multiple output dimensions, the inference model (BO in our cases) needs to consider all the dimensions jointly despite the independence assumption, resulting in one 4-d BO problem instead of three 2-d ones. Having to consider more output dimensions jointly could affect the fitting of each individual dimension, and we intend to explore the scaling performance of `metaGP-gu` on this aspect.

Figure C4 shows the performance comparisons on `syn-mulo`. The x-axis is the number of output dimensions jointly considered in the inference model, which we term as effective output dimension. As references, we include also the performance of other baselines on `syn-mulo`, and note that none of the baselines is affected by the change on the effective output dimensions, including `SLGP` which always takes the maximum of few-shots as scaling factor of input. From the plots, it is clear to see the negative effect of the effective output dimensions on the prediction performance, and the potential explanations are that the inferred scaling on the input would need to compromise between the three outputs and that it is more difficult for BO to explore a larger search space. Therefore, the poorer performance of BO on higher dimensional space can harm the performance of our meta-learning framework.

## Appendix D. The effect of inner-loop gradient update in the evaluations

Here we show the effect of having inner-loop gradient update in various datasets. Judging from Figure D1 and Table D1, the effect of inner-loop gradient update differs according to the datasets and the population models. For example for both the univariate datasets, on `syn-exp` the effects of having gradient update are trivial, while on `semi-C-syn`, a noticeable improvement was observed across all number of shots. And for datasets with multivariate inputs, the inner-loop gradient update seems to be more beneficial in the higher dimensional space. The dataset-dependent improvements are also observed in the real datasets, while gradient update is more helpful in predicting C fiber amplitudes overall, `Mul-B` suffers from having the extra step. Another key factor affecting the performance of the gradient update is the choice of the population model, where on the univariate real datasets, the methods with GP population models are more likely to be affected than those with regression models. Overall in our evaluations, having the inner-loop gradient update shows a positive influence on few-shot performance, and in the cases when the methods actually suffered from the gradient update, the negative effect tends to be small.

## Appendix E. Additional investigations on the effects of population set

We conducted additional investigation targeting `np`. The effect of `np` has a more complicated nature, as it affectes not only the size of data from each subject but indirectly also the normalized population set. Recall that to obtain the population models, the data has to be normalized by its respective maximal values. During subject generation, the semi-synthetic datasets were injected with zero-mean Gaussian noise, so the absolute maximal fiber responses can be larger than the ground truth *Z*_*y*_, meaning the normalization factor on fiber amplitude could commonly be overestimated if `np` is equal to the largest possible value. However, it is undesirable also to keep `np` too small, for example, if `np` is 1, every point in the normalized population set becomes (1, 1). We would hypotheses an intermediate value as an optimal one, and Figure 7(b) suggested that it might be justified, but still uncertain due to the small gaps among the versions in comparison. Therefore, here, we show in Table E1 the mean and standard deviation of the RMSE distance between the fit used to generate all the subjects and the obtained population fit for various combinations of `ns` and `np`.

The results corroborate that the population set with an intermediate value of `np` did have a closer shape to the ground truth. Secondly, we ran the same evaluations of varying `np`, but this time with datasets of `y nm` as 2, 3 and 5. Note that original one has `y nm` as 1. The results are shown in Figure E1, and they show the similar trends as Figure 7(b) that intermediate value performs the best.

**Figure C4:**
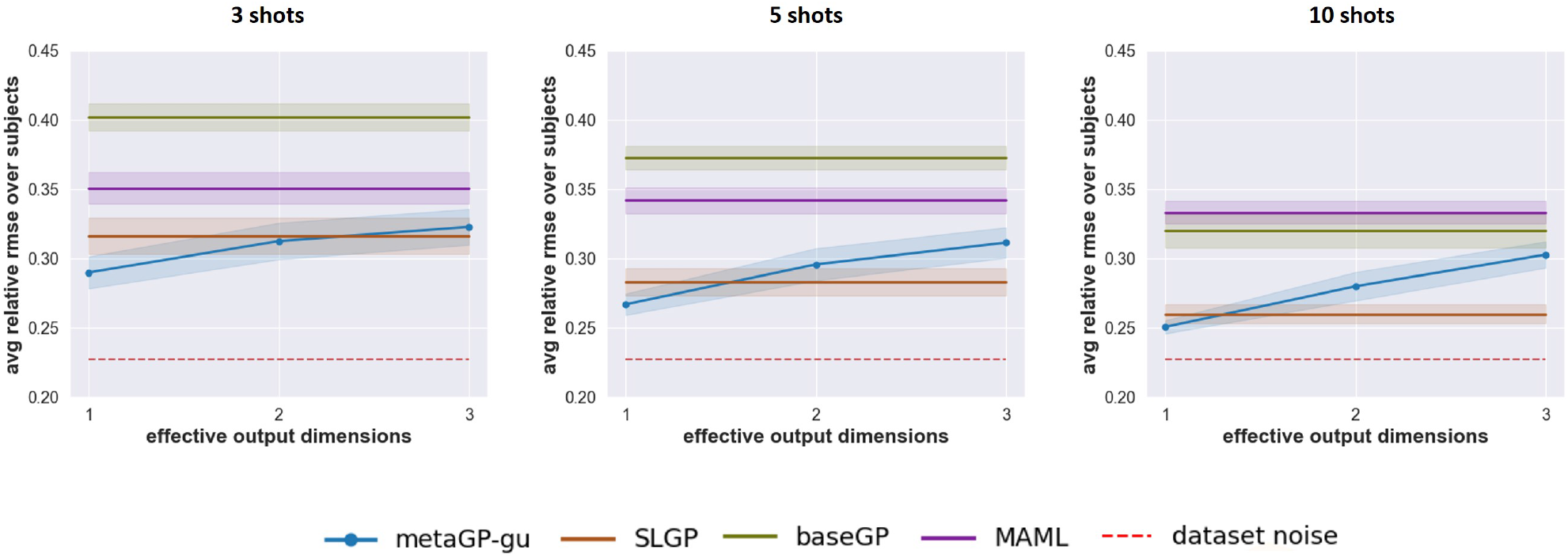
Performance evaluations on scaling in synthetic datasets with multivariate outputs. Plots show performance of all the algorithms on all datasets with effective output dimensions from 1 to 3, with 3, 5 and 10 shots.

**Figure D1:**
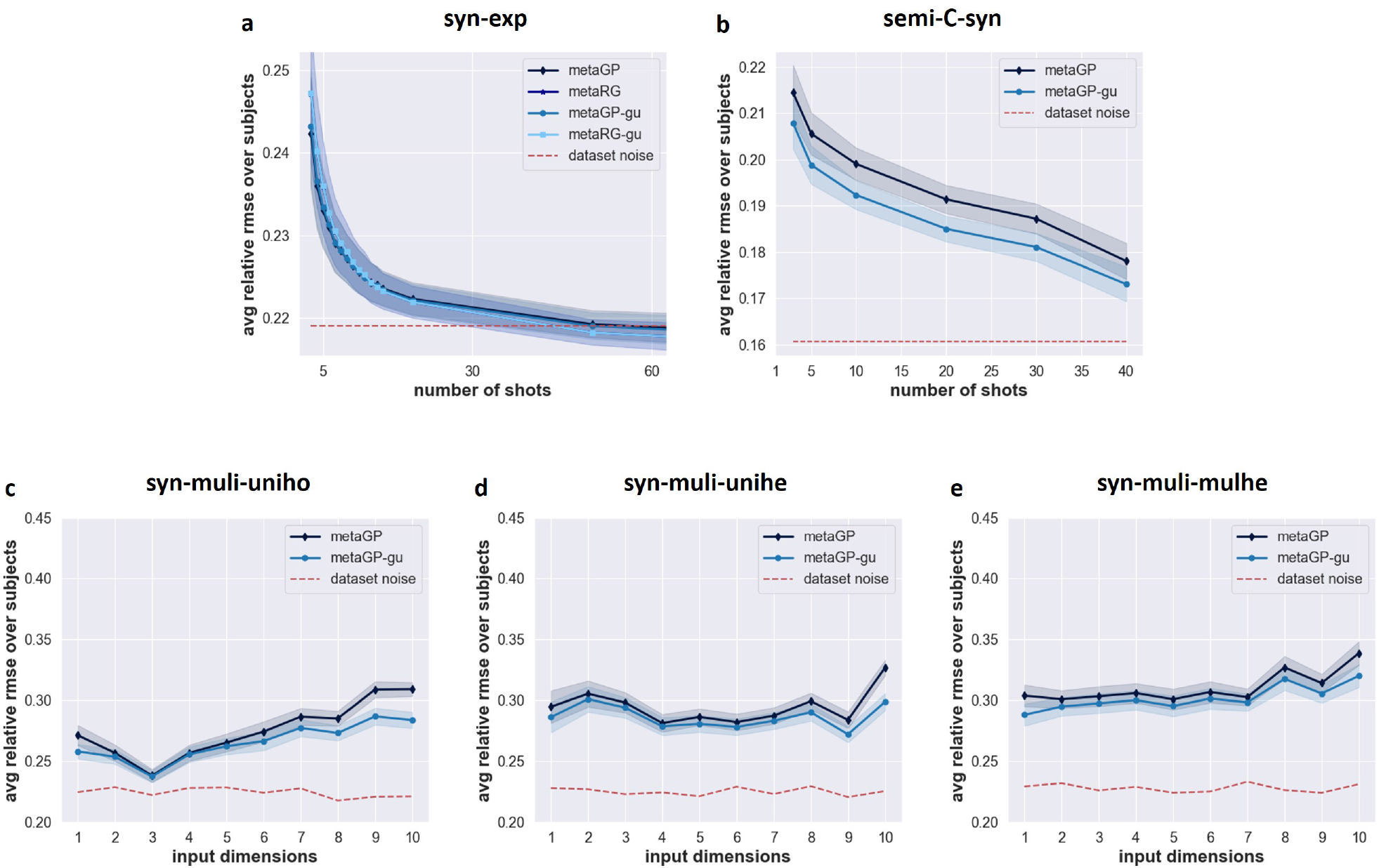
Effect of inner-loop gradient update on five synthetic datasets (a) `syn-exp` (b) `semi-C-syn` (c)

**Table D1:** Effect of inner-loop gradient update on all six regression tasks of the real experimental data. For each algorithm and each task, leave-one-out cross-validation was conducted, and the values reported were its average testing RMSE over all subjects, and over 3 and 5 shots.

|  | EMG-A | chr-B | cBI-C | Mul-A | Mul-B | Mul-C |
| --- | --- | --- | --- | --- | --- | --- |
| metaGP - metaGP-gu | 0.198 | 0.155 | 0.270 | 0.180 | -0.038 | 0.393 |
| metaRG - metaRG-gu | 0.003 | $-3e^{-6}$ | -0.003 | - | - | - |

**Figure E1:**
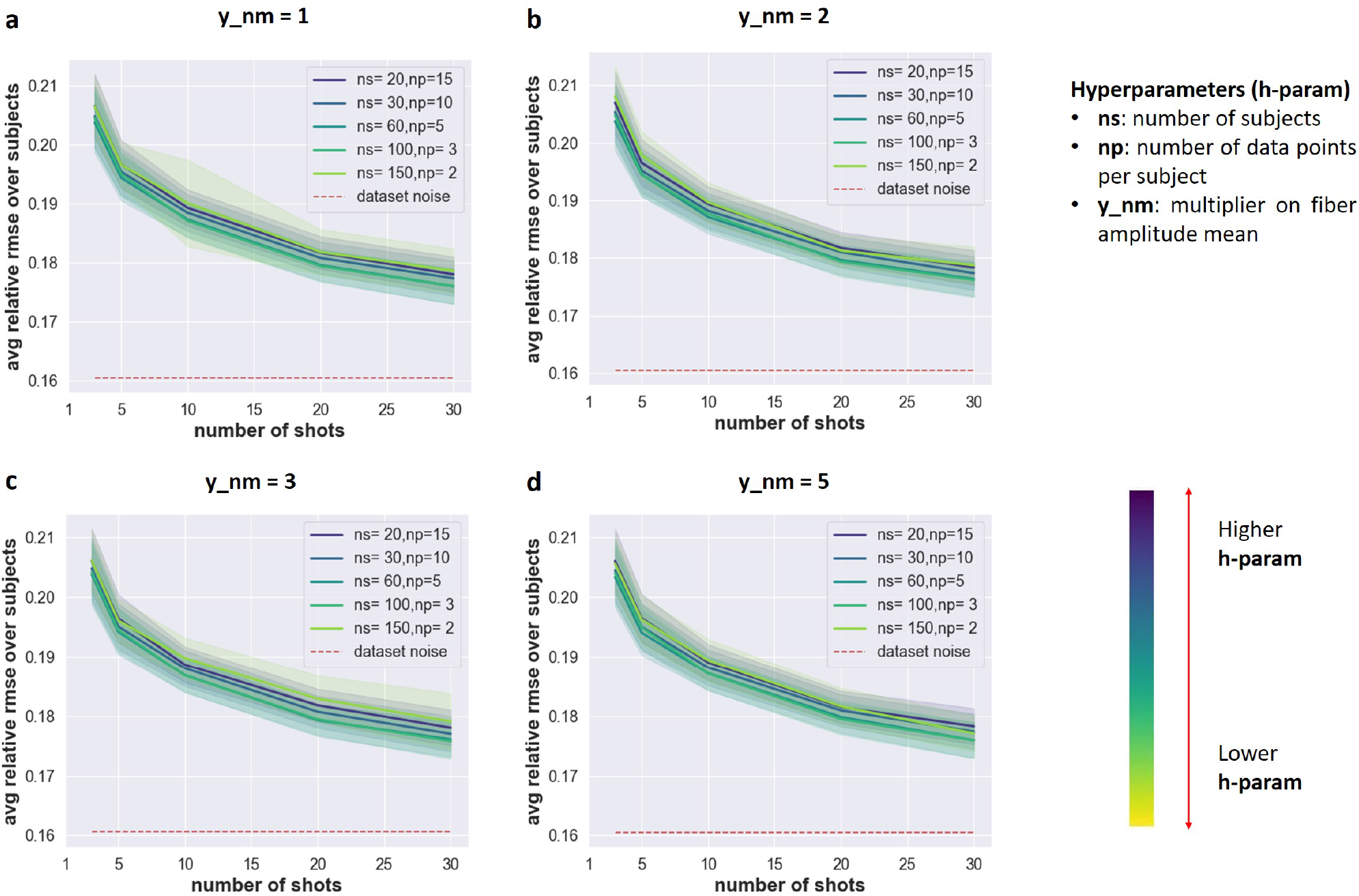
Investigations on the effect of `np` (number of data points per subject) with varying `y nm` (multiplier on ranges of subjects’ fiber amplitude of the population set). Each plots show relative rooted mean squared error: (a) `y nm` as 1, this is exactly the same figure as in Figure 7(b); (b) `y nm` as 2; (c) `y nm` as 3; (d) `y nm` as 5.

**Table E1:**
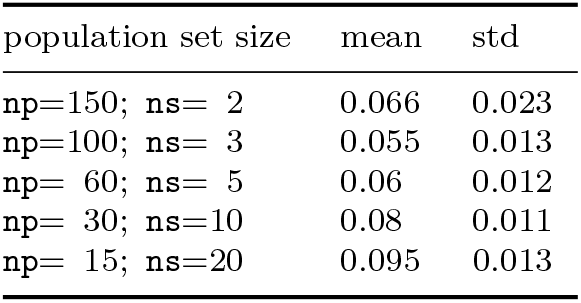
Comparisons among different combinations of the number of subjects and the number of points per subject in the population set. Values reported are the distance between the data generating function and each obtained population fits.

## Reference

[1] Brooker C et al 2021 ECAP-Controlled Closed-Loop Spinal Cord Stimulation Efficacy and Opioid Reduction Over 24-Months: Final Results of the Prospective, Multicenter, Open-Label Avalon Study Pain Practice 21 680–691

[2] Okun M S 2012 Deep-Brain Stimulation for Parkinson’s Disease New England Journal of Medicine 367 1529–1538

[3] Ben-Menachem E 2002 Vagus-nerve stimulation for the treatment of epilepsy The Lancet Neurology 1 477–482

[4] Edwards C A, Kouzani A, Lee K H and Ross R K 2017 Neurostimulation Devices for the Treatment of Neurologic Disorders Mayo Clinic Proceedings 92 1427–1444

[5] Tufail Y, Yoshihiro A, Pati S, Li M M and Tyler W J Tyler 2011 Ultrasonic neuromodulation by brain stimulation with transcranial ultrasound Nat Protoc 6 1453–1470

[6] Handforth A et al 1998 Vagus nerve stimulation therapy for partial-onset seizures: a randomized active-control trial Neurology 51 48–55

[7] Ahmed U, Chang Y C, Zafeiropoulos S, Nassrallah Z, Miller L and Zanos S 2022 Strategies for precision vagus neuromodulation Bioelectronic medicine 8

[8] Wang Y Q, Yao Q M, Kwok J T and Ni L M 2020 Generalizing from a Few Examples: A Survey on Few-shot Learning ACM Comput. Surv. 53 1–34

[9] Goodfellow I, Bengio Y and Courville A 2016 Deep Learning (MIT Press)

[10] Ravi S and Larochelle H 2017 Optimization as a Model for Few-Shot Learning Int. Conf. on Learning Representations

[11] Finn C, Abbeel P and Levine S 2017 Model-Agnostic Meta-Learning for Fast Adaptation of Deep Networks Proc. Int. Conf. on Machine Learning, PMLR 70 1126–1135

[12] Sigrid B, Aleksandra K, Gerhard R and Gregor H 2018 Vagus Nerve as Modulator of the Brain–Gut Axis in Psychiatric and Inflammatory Disorders Front. Psychiatry 9

[13] Rush A J et al. 2005 Vagus nerve stimulation for treatmentresistant depression: a randomized, controlled acute phase trial Biol Psychiatry 58 347–354

[14] George M S, Ward H E Jr, Ninan P T, Pollack M, Nahas Z, Anderson B, Kose S, Howland R H, Goodman W K and Ballenger JC 2008 A pilot study of vagus nerve stimulation (VNS) for treatment-resistant anxiety disorders Brain Stimul. 1 112–121

[15] Merrill C A et al. 2006 Vagus nerve stimulation in patients with Alzheimer’s disease: additional follow-up results of a pilot study through 1 year. J Clin Psychiatr 67 1171–1178

[16] Chakravarthy K, Chaudhry H, Williams K and Christo P J 2015 Review of the Uses of Vagal Nerve Stimulation in Chronic Pain Management Current pain and headache reports 19 54

[17] Koopman F A et al. 2016 Vagus nerve stimulation inhibits cytokine production and attenuates disease severity in rheumatoid arthritis Proc Natl Acad Sci USA 113 8284–8289

[18] Labiner D M and Ahern G L 2007 Vagus nerve stimulation therapy in depression and epilepsy: therapeutic parameter settings Acta Neurol Scand. 115 23–33

[19] Musselman E D, Pelot N A and Grill W M 2019 Empirically Based Guidelines for Selecting Vagus Nerve Stimulation Parameters in Epilepsy and Heart Failure Cold Spring Harb Perspect Med. 9

[20] He S M, Teagle H F B and Buchman C A 2017 The Electrically Evoked Compound Action Potential: From Laboratory to Clinic Front. Neurosci. 11

[21] Chang Y C et al. 2020 Quantitative estimation of nerve fiber engagement by vagus nerve stimulation using physiological markers Brain Stimul. 13 1617–1630

[22] McAllen R M, Shafton A D, Bratton B O, Trevaks D and Furness J B 2018 Calibration of thresholds for functional engagement of vagal A, B and C fiber groups in vivo Bioelectronics in medicine 1 21–27

[23] Qing K Y, Wasilczuk K M, Ward M P, Phillips E H, Vlachos P P, Goergen C J and Irazoqui P P 2018 B fibers are the best predictors of cardiac activity during Vagus nerve stimulation Bioelectronic Medicine 4 5

[24] Rasmussen C E and Williams C K I 2006 Gaussian Processes for Machine Learning (MIT Press)

[25] Wang Z, Gehring C, Kohli P and Jegelka S 2018 Batched Large-scale Bayesian Optimization in High-dimensional Spaces International Conference on Artificial Intelligence and Statistics

[26] Sun Q, Liu Y, Chua T and Schiele B 2019 Meta-Transfer Learning for Few-Shot Learning 2019 IEEE/CVF Conference on Computer Vision and Pattern Recognition 403-412 doi: 10.1109/CVPR.2019.00049

[27] Perez E, Strub F, de Vries H, Dumoulin V and Courville A 2018 FiLM: Visual Reasoning with a General Conditioning Layer Proceedings of the AAAI Conference on Artificial Intelligence 32

[28] Bengio Y, Courville A and Vincent P 2013 Representation Learning: A Review and New Perspectives IEEE Transactions on Pattern Analysis and Machine Intelligence 35 1798–1828

[29] Vuorio R, Sun S H, Hu H X and Lim J J 2019 Multimodal Model-Agnostic Meta-Learning via Task-Aware Modulation 2019 Advances in Neural Information Processing Systems

[30] Zhang Y and Yang Q 2021 A Survey on Multi-Task Learning IEEE Transactions on Knowledge and Data Engineering

[31] Sutton R S and Barto A G 2018 Reinforcement learning: An introduction (MIT press)

[32] Bertsekas D P 2000 Dynamic Programming and Optimal Control (Athena Scientific)

[33] Bishop C M 2007 Pattern Recognition and Machine Learning (Springer New York)

[34] Wilson A G, Hu Z, Salakhutdinov R and Xing EP 2016 Deep Kernel Learning. Proc. Int. Conf. on Machine Learning, PMLR 51 370–378

[35] Brochu E, Cora V and Freitas N E 2010 A Tutorial on Bayesian Optimization of Expensive Cost Functions, with Application to Active User Modeling and Hierarchical Reinforcement Learning ArXiv:abs/1012.2599

[36] Mockus J, Tiesis V and Zilinskas A 1978 The Application of Bayesian Methods for Seeking the Extremum Towards Global Optimization 2 117–129

[37] Paszke A et al PyTorch: An Imperative Style, High-Performance Deep Learning Library 2019 Advances in Neural Information Processing Systems 32 8024–8035

[38] Balandat M, Karrer B, Jiang D R, Daulton S, Letham B, Wilson A G and Bakshy E 2020 BoTorch: A Framework for Efficient Monte-Carlo Bayesian Optimization Advances in Neural Information Processing Systems 33

[39] Gardner J R, Pleiss G, Bindel D. Weinberger K Q and Wilson A G 2018 GPyTorch: Blackbox Matrix-Matrix Gaussian Process Inference with GPU Acceleration Advances in Neural Information Processing Systems 31

[40] Deleu T, Würfl T, Samiei M, Cohen J P and Bengio Y 2019 Torchmeta: A Meta-Learning library for PyTorch ArXiv:abs/1909.06576

[41] Virtanen P et al 2020 SciPy 1.0: Fundamental Algorithms for Scientific Computing in Python Nature Methods 17 261–272

[42] Kallewaard J W et al. 2021 Real-World Outcomes Using a Spinal Cord Stimulation Device Capable of Combination Therapy for Chronic Pain: A European, Multicenter Experience J. Clin. Med. 10

[43] van Bueren N E R, Reed T L, Nguyen V, Sheffield J G, van der Ven S H G, Osborne M A, Kroesbergen E Rapidly Inferring Personalized Neurostimulation Parameters with Meta-Learning 20

[44] H Kadosh R C 2021 Personalized brain stimulation for effective neurointervention across participants PLOS Computational Biology 17 1–24

[45] Hollunder B, Rajamani N, Siddiqi S H, Finke C, Kühn A A, Mayberg H S, Fox M D, Neudorfer C and Horn A 2022 Toward personalized medicine in connectomic deep brain stimulation. Progress in Neurobiology 210

[46] Razavi B et al 2020 Real-world experience with direct brain-responsive neurostimulation for focal onset seizures. Epilepsia 61 1749–1757

[47] Picillo M, Lozano A M, Kou N, Puppi Munhoz R and Fasano A 2016 Programming Deep Brain Stimulation for Parkinson’s Disease: The Toronto Western Hospital Algorithms. Brain stimulation 9 425–437

[48] Brown T B et al. 2020 Language Models are Few-Shot Learners Advances in Neural Information Processing Systems 33

[49] Finn C, Yu T, Zhang T, Abbeel P and Levine S 2017 One-Shot Visual Imitation Learning via Meta-Learning Proceedings of the 1st Annual Conference on Robot Learning, PMLR 78 357–368

[50] Altae-Tran H, Ramsundar B, Pappu A S and Pande V 2017 Low Data Drug Discovery with One-Shot Learning ACS Central Science 3 283–293

[51] Hospedales T, Antoniou A, Micaelli P and Storkey A 2021 Meta-Learning in Neural Networks: A Survey IEEE Transactions on Pattern Analysis and Machine Intelligence 1

[52] Vinyals O, Blundell C, Lillicrap T, Kavukcuoglu K, and Wierstra D 2016 Matching networks for one shot learning Advances in Neural Information Processing System 29

[53] Santoro A, Bartunov S, Botvinick M, Wierstra D and Lillicrap T 2016 Meta-learning with memory-augmented neural networks Proc. Int. Conf. on Machine Learning, PMLR 48 1842–1850

[54] Blei D, Ng A, and Jordan M 2003 Latent dirichlet allocation. Jounral of Machine Learning Research 3 993–1022

[55] Grant E, Finn C, Levine S, Darrell T. and Griffiths T L 2018 Recasting Gradient-Based Meta-Learning as Hierarchical Bayes Int. Conf. on Learning Representations

[56] Amit R and Meir R 2018 Meta-Learning by Adjusting Priors Based on Extended PAC-Bayes Theory Proc. Int. Conf. on Machine Learning, PMLR

[57] Kingma D P and Ba J 2015 Adam: A Method for Stochastic Optimization Int. Conf. on Learning Representations Rapidly Inferring Personalized Neurostimulation Parameters with Meta-Learning

